# Human Digital Twin: Automated Cell Type Distance Computation and 3D Atlas Construction in Multiplexed Skin Biopsies

**DOI:** 10.1101/2022.03.30.486438

**Authors:** Soumya Ghose, Yingnan Ju, Elizabeth McDonough, Jonhan Ho, Arivarasan Karunamurthy, Chrystal Chadwick, Sanghee Cho, Rachel Rose, Alex Corwin, Christine Surrette, Jessica Martinez, Eric Williams, Anup Sood, Yousef Al-Kofahi, Louis D. Falo, Katy Börner, Fiona Ginty

## Abstract

Mapping the human body at single cell resolution in three-dimensions (3D) is an important step toward a “digital twin” model that captures important structure and dynamics of cell-cell interactions. Current 3D imaging methods suffer from low resolution and are limited in their ability to distinguish cell types and their spatial relationships. We present a novel 3D workflow: MATRICS-A (<u>M</u>ultiplexed Im<u>a</u>ge <u>T</u>hree-D <u>R</u>econstruction and <u>I</u>ntegrated <u>C</u>ell <u>S</u>patial - <u>A</u>nalysis) that generates a 3D map of cells from multiplexed images and calculates cell type distance from endothelial cells and other features of interest. We applied this workflow to multiplexed data from sequential skin sections from younger and older donors (n=10; 33-72 years) with biopsies from ten anatomical regions with different sun exposure effects (mild, moderate-marked). Up to 26 sequential sections from each sample underwent multiplexed imaging with 18 biomarkers covering 12 cell types (keratinocytes (granular, spinous, basal), epithelial and myoepithelial cells, fibroblasts, macrophages, T helpers, T killers, T regs, neurons and endothelial cells, markers of DNA damage and repair (p53, DDB2) and cell proliferation (Ki67). Following cell classification, the tissue and classified cells were reconstructed into 3D volumes. A significant inverse correlation between DDB2 positive cells and age was found (corr= -0.78, adj. p=0.047). This suggests reduced capacity for repair in non-cancer older sun-exposed individuals. While absolute immune cell count did not differ by age or sun exposure, the ratio of T Helper/T Killer cells was positively correlated with age (corr=0.82, adj. p=0.048) This is the first such 3D study in skin and paves the way for cataloging more cell types and spatial relationships in aging and disease in skin and other organs.

## Introduction

The National Institutes of Health’s (NIH) Human Biomolecular Atlas Program (HuBMAP) aims to create a comprehensive high-resolution atlas of all cells in the healthy human body using data from multiple laboratories across the US and Europe^1^. Integrating and harmonizing the data derived from these samples and “mapping” them into a common three-dimensional (3D) space is a major challenge. HuBMAP, in close collaboration with 16 other international consortia and projects, is systematically constructing a Human Reference Atlas^2^. At the core of this Atlas is a common coordinate framework (CCF) that supports spatially and semantically explicit human tissue registration and exploration. The completed Atlas will support the design of a “digital twin” for healthy men and women that can be parameterized in support of precision health and medicine. The CCF has two key components: (1) anatomical structures, cell types, and biomarkers (ASCT+B) tables that name key entities and link them to existing ontologies (e.g., Uberon multi-species anatomy ontology, Foundational Model of Anatomy Ontology [FMA], Cell Ontology [CL], or HUGO Gene Nomenclature [HGNC]) and (2) a 3D reference object library that spatially defines the 2D and 3D structures of anatomical structures and cell types, and characterizes their spatial relationships. Specifically, this paper computes distance distributions for immune and other cell types to the nearest blood vessel in three dimensions for 3D digital skin biopsy data.

Skin is the largest organ and is composed of at least 36 different cell types (documented in version 1.1 of ASCT+B^3^) and a vast microenvironment of over 16 anatomical structures—including glandular structures, hair follicles, vasculature, and immune system components. At least 70 protein biomarkers are needed to identify these cell types and anatomical structures^3^, and even more if increased cellular granularity and functionality are needed. While several single cell studies or atlases of human skin have been conducted in recent years^4,5^, these have focused on single cell RNAseq analysis and do not focus on 2D *in situ* or 3D spatial analysis of cell types and proteins. To that end, we have developed a new workflow for 3D reconstruction of multiplexed skin cell types and cell distance distributions (MATRICS-A). The workflow enabled 3D evaluation of aging and sun exposure effects on the epidermis and dermis, including epidermal localization of ultraviolet (UV) radiation damaged cells (e.g., p53 mutations), DNA repair (DDB2), proliferation (Ki67), and immune cell counts and spatial distances to endothelial cells. While much work has been done on characterizing precancerous and cancerous skin, there is less understanding of cellular changes in otherwise healthy individuals across the lifecycle. UV is a major environmental stressor, with the risk of developing disease substantially increasing with age and exposure ^6–8^.

The novelty of our approach lies in the reconstruction of a 3D volume using multiplexed images and multiple cell types and calculation of spatial distances. Compared to previous reconstruction methods^9–12^, our approach allows more precise and biologically relevant analysis of cellular relationships and has broad applications in a disease context where immune response, angiogenesis and microenvironment interrelationships provide important mechanistic insights into disease progression and tumor heterogeneity^10^. **Figure 1:** illustrates the end-to-end process undertaken for this analysis. **Fig 1A:** Healthy skin biopsies were embedded into a single formalin-fixed and paraffin-embedded (FFPE) tissue block. We used the human male and female skin 3D reference organ to spatially register and semantically annotate the biopsies via the HuBMAP Registration User Interface^13^; **Fig. 1B:** Skin biomarkers were identified using the skin ASCT+B tables and corresponding antibodies were validated^3^; **Fig. 1C:** A block with 12 samples underwent micro CT imaging and was then sectioned into 26 serial sections for highly multiplexed immunofluorescence imaging using 18 protein and cell type markers. **Fig. 1D:** Cell classification was conducted for each section using a hybrid supervised (deep learning-based) and unsupervised (probability-based) workflow. Serial sections and cell types then underwent 3D reconstruction. Next, the 3D spatial location of immune cells was used to compute cell distance distributions from endothelial cells for each age group and sun exposure effect. Differences in epithelial composition of p53, Ki67, and DDB2 positive cells and their distance from the skin surface were evaluated to assess the effects of sun exposure on skin cells.

**Figure 1:**
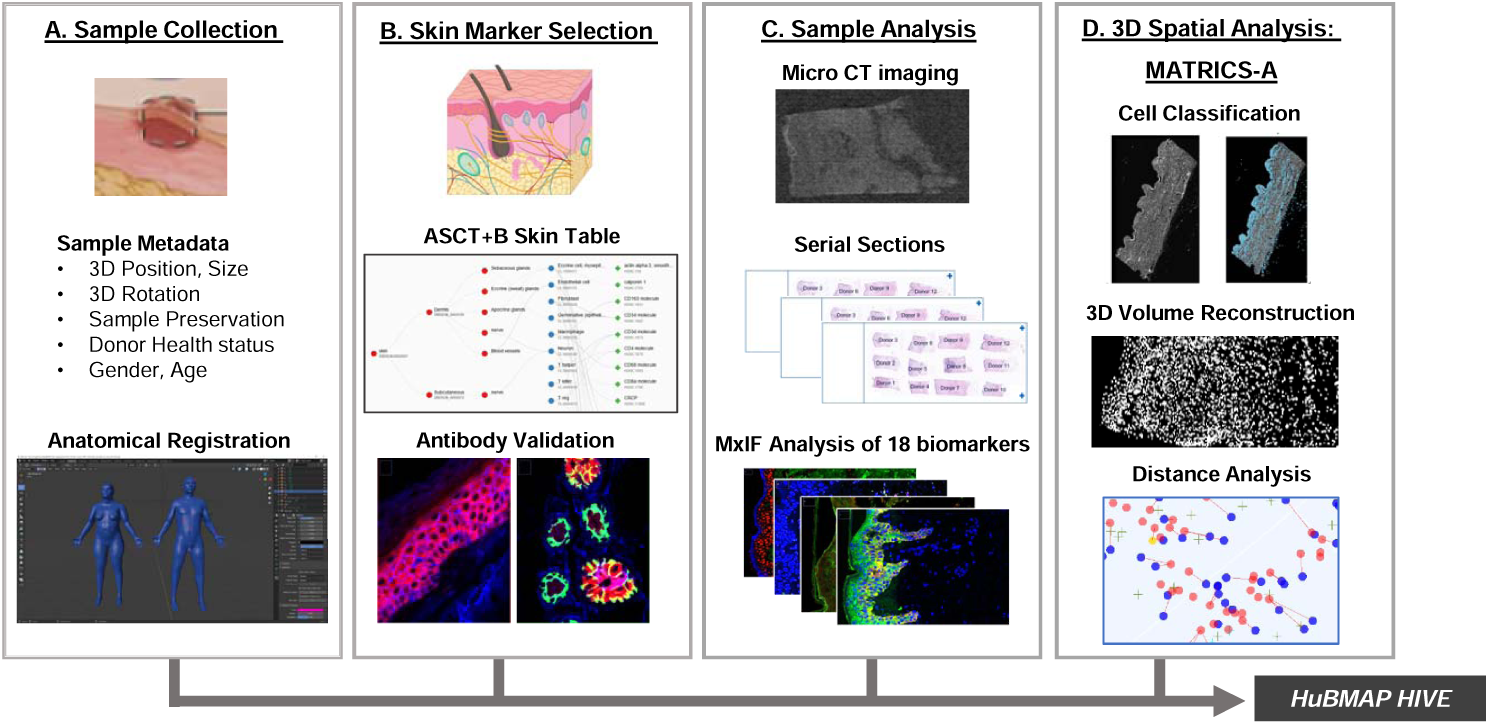
End-to-end workflow for generation of a 3D skin map of cell types and distances

## Results

### Skin Anatomical Structure and Cell Type + Biomarker (ASCT+B) Table and 3D Reference Organs

For this work, an ASCT+B table as well as a male and female 3D reference organ were constructed for skin.

The 3D skin reference organs^14,15^ are shown in **Fig. 1A and Supplementary Fig. S2**. They were derived from the National Library of Medicine (NLM) Visible Human project^16^ data. The male and female reference organs were added to the HuBMAP Registration User Interface ^13^ making it possible to formally register all biopsy tissue samples into the evolving Human Reference Atlas (HRA). The result is metadata for each tissue block that documents the size, spatial location, and rotation in 3D. All registered tissue blocks used in this study can be interactively explored in the HuBMAP portal’s CCF Exploration User Interface (CCF-EUI)^17^. The version 1.1 skin ASCT+B master table^3^ achieves three critical elements: (1) capturing the *part_of* relationships between 15 anatomical structures that are linked to their respective Uberon IDs, (2) featuring the 36 skin cell types (linked to CL) that are *located_in* one or more of these anatomical structures, and (3) identifying 70 protein biomarkers (linked to HGNC ontology) that are commonly used to *characterize* the 36 cell types. This paper focuses on a subset of the entities in the skin master table, namely 18 biomarkers (including the functional markers Ki67, p53 and DDB2) and 12 cell types. A sample of the skin ASCT+B reporter is shown **Fig. 1B and Supp. Fig. S4** and a complete view of all 36 cell types and 70 biomarkers is accessible at our Companion Website^18^.

### Hybrid supervised and unsupervised approach for precise cell classification

To address specific questions around aging and UV damage, we focused on segmentation of epithelial, immune, and endothelial cells using an automated, hybrid unsupervised and supervised framework (See **Methods** and **Supp. Figs. S6 and S7**). Given the amount of data required for 3D reconstruction of multiplexed images, it would have been a cumbersome and time-consuming exercise to design this study with only deep learning (DL) models. Typically, a large number of manually annotated cells are required to develop a deep learning-based segmentation model and manual annotation introduces inter and intra-rater variabilities. To achieve our goal, we adopted a hybrid supervised and unsupervised model where a supervised DL model was used for DAPI segmentation and unsupervised Gaussian mixture models (GMM) were used for probabilistic segmentation of individual biomarkers. The union of probabilistic biomarker segmentation with DL based nuclei segmentation resulted in a robust cell classification model with high sensitivity, specificity, and accuracy for all markers from significantly less manually annotated data than traditional deep learning models^19^ (**Figure 2B-E**). Gaussian mixture models (GMM) were used for probabilistic segmentation of epithelium (cytokeratin) and cells undergoing DNA damage (p53), repair (DDB2), and proliferation (Ki67). A similar probabilistic segmentation workflow was used for endothelial and immune cell classification.

**Figure 2:**
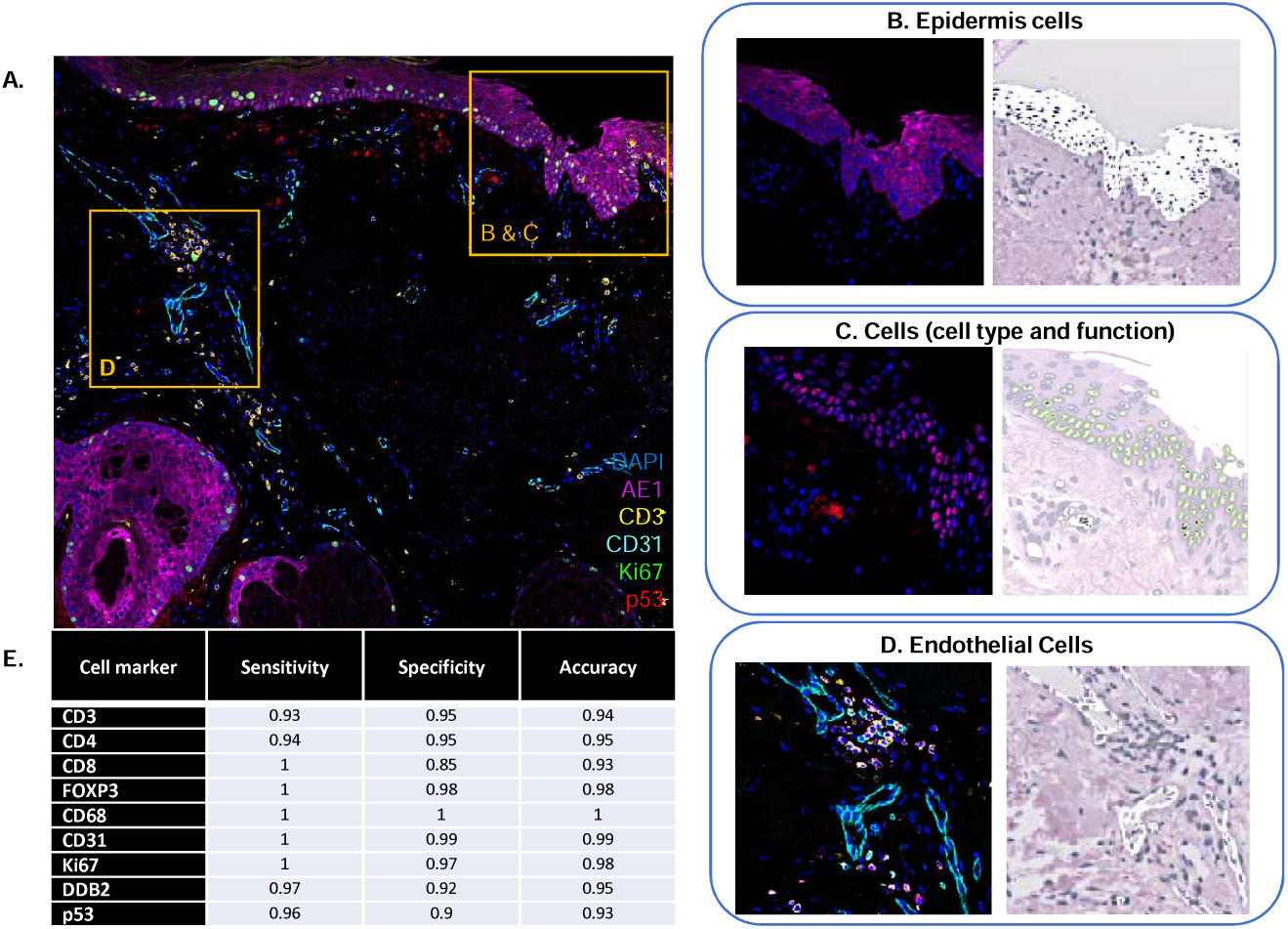
Skin cell classification and performance

### 3D skin volume and cell reconstruction provide a novel framework for spatial analysis

Our 3D reconstruction framework allowed us to create 3D volumes from autofluorescence (AF) images of multiplexed serial sections and spatially map cell types in 3D for further analysis (See **Fig. 3** and **Methods)**. We achieved a mean dice similarity coefficient (DSC)^21^ or overlap accuracy of 0.95+/-0.04 for 24 serial sections for 10 volumes after registration (the last 2 sections were used for deep learning training and excluded). An overlap DSC of 0.90 is classified as high quality in the 3D registration or reconstruction community^9^. We further computed normalized cross correlation between the serial sections for all samples to quantify registration quality and found 0.6+/-0.07 for all AF serial sections, which is comparable to established 3D reconstruction methods^9^ in the community considering large scale deformation and tissue damage during the course of cyclic multiplexing. Compared to traditional registration approaches^23^ used for 3D reconstruction, we automatically segment out the AF image to create a mask, which focuses registration in the region of interest (ROI) and filters out background noise that may interfere with the registration process. This approach improves our inter-slice registration accuracy and speed. Further, compared to landmark-constrained 3D histological imaging^24^, minimal manual intervention is required for an accurate registration and reconstruction of the 3D volume. Within this workflow, a single slide from the entire volume is identified manually based on tissue quality as the reference image; the rest of the process for registration and 3D reconstruction is completely automatic. Compared to image similarity-based alignment^9,22^, automatic block correspondences are used for initial affine alignment. This approach improves registration accuracy especially in scenarios where there is tissue damage, tissue deformation, imaging artifacts and noise. Use of block correspondences improves registration speed compared to image similarity-based alignment methods^22,23^. To the best of our knowledge, this is the first time a 3D reconstruction method has been applied to whole slide multiplexed images - allowing spatial mapping of distributions of multiple cell types in 3D without sacrificing image resolution and the number of biomarkers that could be targeted, compared to other 3D or reconstruction methods. Registering the 3D reconstructed volumes to micro CT images during registration is valuable when there is deformation and/or wear and tear in the tissue samples during the cyclic staining process. The co-registration also allows us to map microfeatures (e.g., cell types) to macro imaging features and opens future possibilities for application to other organs and their disease states and progression. We recognize that there may be instances where CT is not available and as such, we designed our 3D reconstruction model to be flexible and able to perform reconstruction in the absence of the CT images.

**Figure 3:**
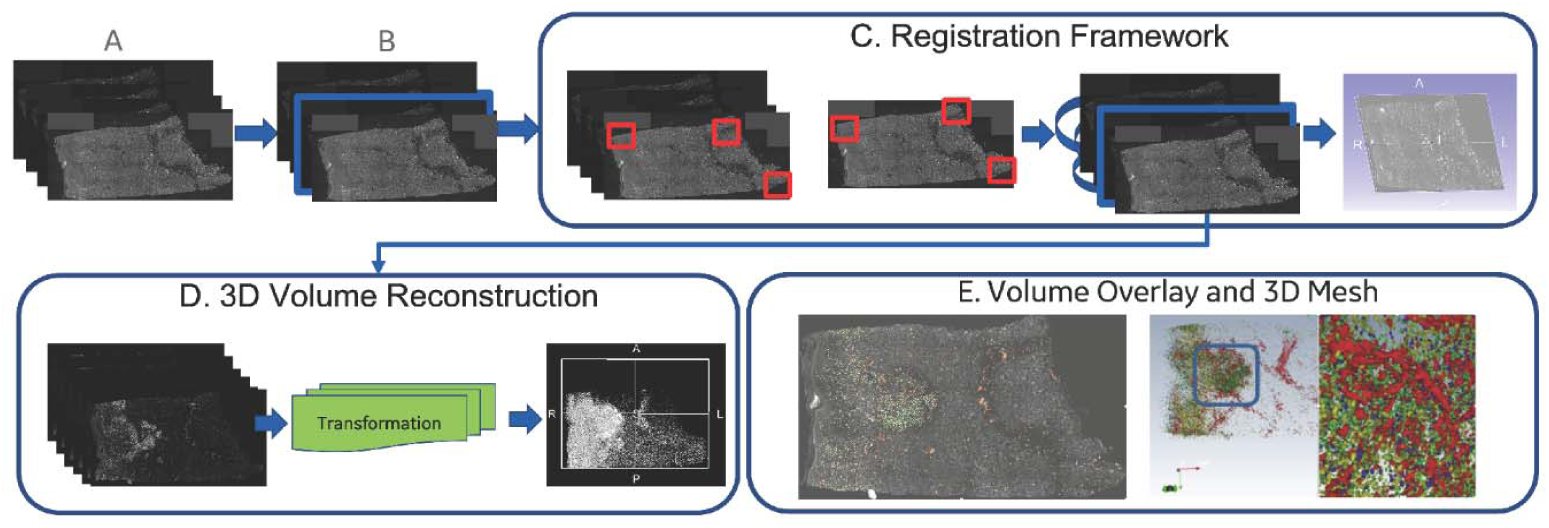
3D Skin Volume Reconstruction Workflow. **3A and 3B** – A reference autofluorescence image from the sequentially imaged sections is used to initiate registration; **C)** A patch-based local correspondences in 2D serial sections was used for affine registration followed by deformable registration to account for tissue deformation; **D)** 3D reconstruction of the biomarkers are achieved by mapping of the biomarkers in 3D using the affine and deformable transformation map and refinement is achieved by registration to micro CT image; **E)** 3D volumes of biomarkers are overlaid on AF volume and 3D mesh of individual biomarkers generated.

### Evidence for reduced DNA repair capacity with aging

DDB2 (damage specific DNA binding protein 2) is the smaller subunit of a heterodimeric protein complex (DDB1 and DDB2) that participates in nucleotide excision repair (NER), which is the principal pathway for countering cytotoxic and mutagenic effects of UV-R^25^. Cell levels of DDB2 are essential for stabilizing DDB1 on damaged DNA^26^ and mutations in the DDB gene lead to xeroderma pigmentosum (XP) complementation group E, which is characterized by increased sensitivity to UV light and a high predisposition for skin cancer^27^. Tumor protein P53 (p53) is activated to counter the DNA damage arising from UV irradiation and a steady low level of UV exposure could lead to continuous^28^ and over expression of wild type or mutant p53, resulting in its accumulation in the nucleus^29^. Positive p53 staining in younger patients has been shown to be limited to the epidermis, increasing progressively with age where it extends deeper into the hair follicles and glands in older patients^28^. While a significant relationship between p53 positive cells and age and sun exposure was not found (**Supp. Fig. S8B**), there was a significant inverse relationship between DDB2 and age (corr= -0.78, adjusted p-value (adj. p) =0.05). This suggests decreased capacity for DNA repair with aging. Although a significant difference was not found with sun exposure, the high correlation between age and sun exposure effects does not rule out further effects of sun damage on DNA repair capacity. The ratio of DDB2/p53 positive cells showed a similar inverse trend with age (corr = -0.59, adj. p=0.16), suggesting lower repair/higher mutation rate in older donors. Ki67 cell count in epidermis region was not significantly correlated with age or sun exposure. This agrees with other studies that have shown that Ki67 expression levels vary in different anatomical locations of skin and do not increase until actinic keratosis (damage to keratinocytes in the lower third of the epidermal layer) or a pre-cancerous/cancerous (full epidermis thickness) *in situ* state is reached^30–32^. Distance to the cell surface was also measured for p53 and Ki67 (higher counts closer to the skin surface is an indicator for early pre-cancerous lesions) and DDB2 (**Figure 4A**). No significant differences were found in relation to aging or sun exposure and distance of these cells to the skin surface. In three cases (region 2, 4 and 9), there was a very wide distribution of values (0-1500 µm from the skin surface), which was due to the presence of a hair follicle in each case (**Figure 54A**). In all other cases, the distance ranged between 0-500 µm.

**Figure 4A:**
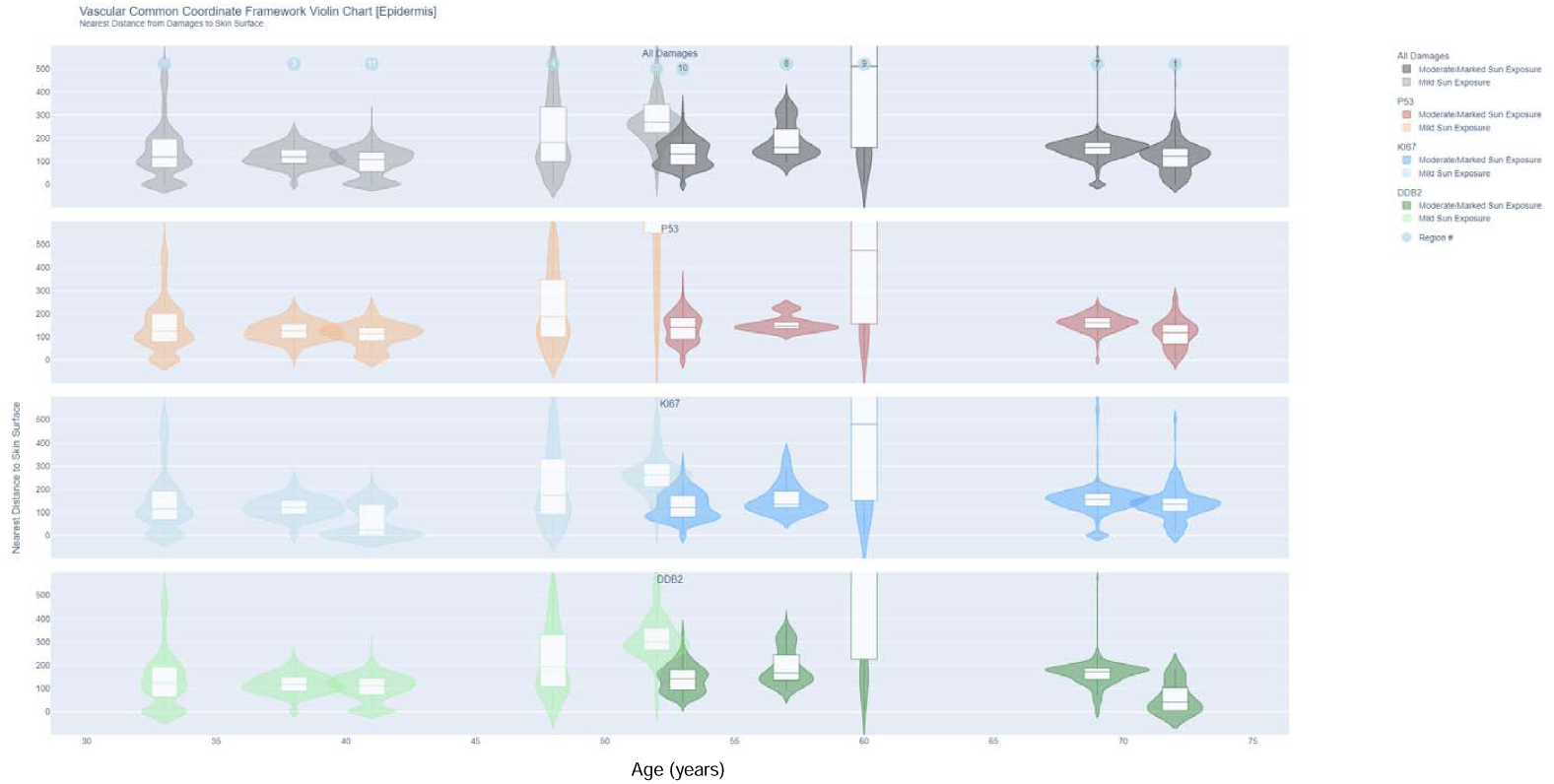
Violin plots of epidermis cells positive for DDB2, p53 and Ki67 and distance to skin surface

**Figure 4B:**
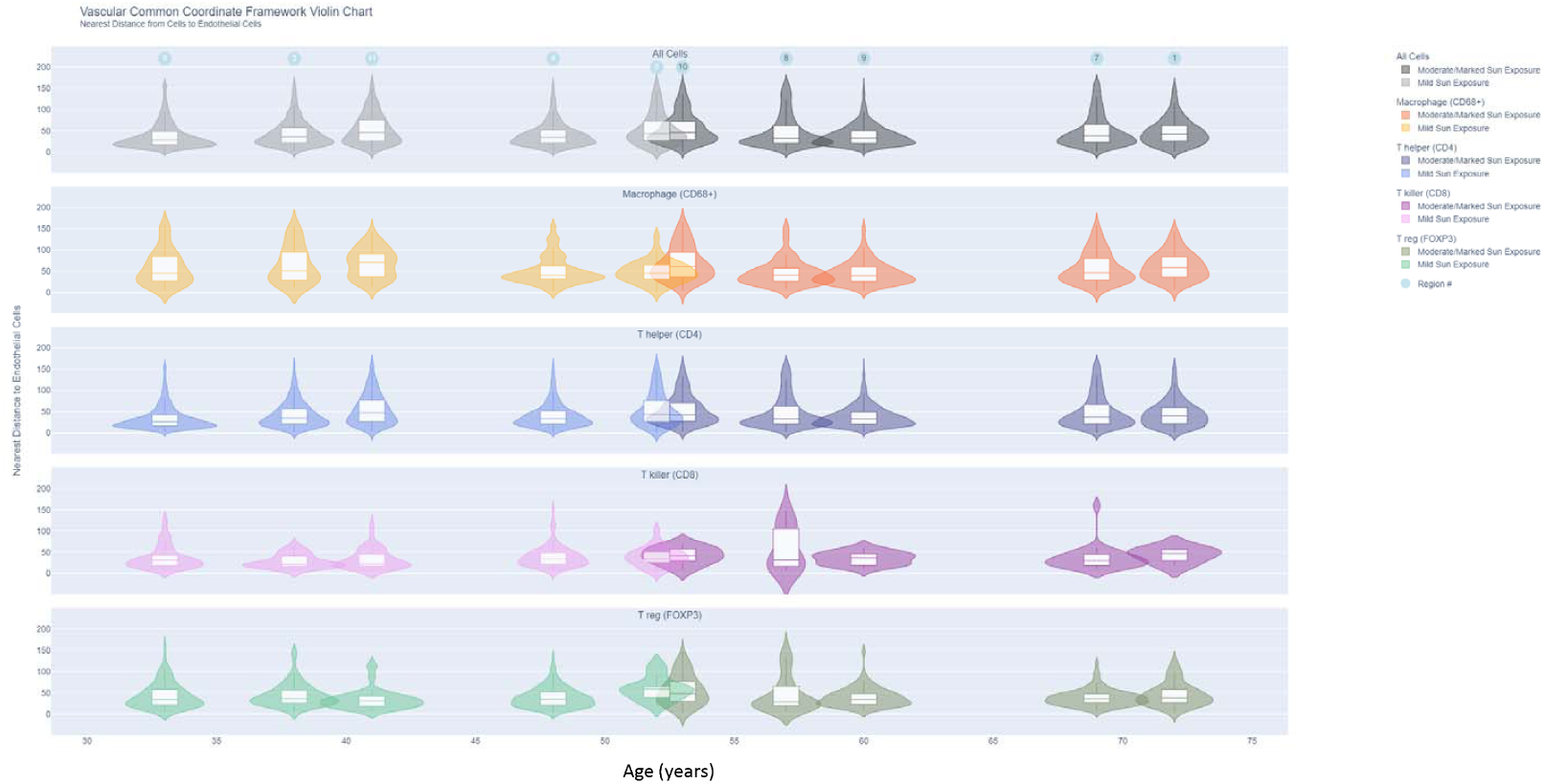
Violin plots of distance of immune cells from nearest endothelial cell by age and exposure

### No change in counts of immune and endothelial cells with age and sun exposure effects

Skin is a huge reservoir of immune cells, with approximately 20 billion T cells across its entire surface area, as estimated by Clark *et al*^33^. It has been demonstrated that in normal skin, more than 95% of those T cells are memory or helper T cells^34^ with T reg cells estimated to be 10% of total population^35^. We found very similar profile in our cohort (T helper: 89 ± 5%; T killer: 2 ± 5%; T reg: 9 ± 5%). Clark *et al*. quantified 590 T cells in a 1 cm (width) × 5-μm section of normal sun-exposed skin (SD = 105, *n* = 25) and a mean of 520 T cells in normal sun-protected skin (SD = 245, *n* = 11) with no difference found between sun exposed/protected skin. Based on their estimation of T cells per cm^2,^ (equivalent to 2 × 10^3^ sections that are 1 cm wide × 5 μm thick), they estimated ∼1 million T cells resident in each cm^2^ of normal skin. Using the entire surface area for a typical 70 kg male (1.8 m^2^), they estimated 1.96 × 10^10^ to 2.12 × 10^10^ total skin resident T cells. Replicating this calculation, we selected a representative whole image for each of the 10 donors and used the median of T cell counts for all images to derive a representative number. We quantified 712 T cells in a 1 cm (width) x 5-μm section (SD=329, n=10) with 685 T cells in mild sun-exposed skin (SD=321, n=5) and 739 T cells in moderate-marked sun exposed skin (SD=337, n=5), with no significant difference between the two groups. Similar to Clark *et al*., we calculated ∼1.4 million T cells in 1 cm^2^⍰ of normal skin and 2.5 × 10^10^⍰ total skin resident T cells across the entire surface area of 1.8 cm^2^. Using the 3D reconstructed volumes from the serial sections of whole slide images, we calculated T cell count after normalizing for sample volume. Average T cells/cm^3^ skin was 33,545,428 (SD 15,636,210) with 28,673,399 (SD=10,670,368) in mild sun exposure samples and 38,417,457 (SD= 19,414,079) with marked exposure (NS). Overall, there were no significant differences in normalized counts (adjusted for tissue volume) in macrophages, T killer cells, T helper cells, or T reg cells by age or in donors with mild vs. moderate-marked sun exposure. A significant positive relationship between T helper/T killer ratio and age was found (corr=0.82, adj. p=0.048). Notably, one donor with marked sun exposure and rheumatoid arthritis (region 1, age 72 years) had the lowest T helper and T killer cell count (**Supp. Fig. S9**) compared to other donors.

### Immune cells and distance from endothelial cells in the dermis

Constructing a vasculature-based coordinate system makes sense biologically as almost every living cell must be within a small distance to a blood vessel (100 µm to 1 mm, depending on the tissue) in order to receive oxygen^36^. The skin’s vasculature, found in its dermal layer, is responsible for temperature regulation, the diffusion of immune cells and nutrient-rich plasma, and barrier loss of body fluid^37^. Aging has been shown to reduce the size and density of blood and lymphatic vessels in the skin as well as disrupt its structure^38^. Chung et al observed an inverse relationship of dermal blood vessel numbers and size with age in sun-damaged, but not in sun-protected skin^39^, with intrinsically aged skin characterized by a reduction in vessel size alone. In this study, we found no significant differences in endothelial cell numbers, regardless of age or sun exposure, however since we did not measure vessel size, we cannot rule out the possibility that there were changes in vessel size. Using the 3D reconstructed data, we also computed the distance of T reg cells, T helper cells, T killer cells and macrophages to nearest endothelial cells. Distance distributions to vasculature cells grouped by age and sun exposure are shown as violin plots in **Fig. 4B and Supp. Fig S10**. An inverse correlation with weak significance between T killer cell count within 100 µm of endothelial cells (normalized by total endothelial cell counts) and age was found (corr= -0.73, adjusted p=0.08). The implications of this are unclear without further validation in a larger group of subjects.

### Interactive visualization of 3D reconstructed skin volumes

Understanding and communicating the 3D spatial location and distance relationships of multiple cell types and supporting the comparison of cell type distance distributions across donors and conditions is non-trivial. For this study, two interactive visualizations were developed to serve this need: (1) a 3D vasculature CCF visualization (VCCF) and (2) interactive cell type distance distribution violin plots. The 3D VCCF Visualization tool makes it possible to examine one 3D reconstructed tissue block at a time. Users can view one or more serial sections; they can view one or more cell types/markers in these sections; they can review the automatically updated distance distribution plots below the 3D skin visualization; plus, they can access the virtual H&E dataset for histological context. Two views are provided: (1) distance to skin surface, focusing on the composition of p53, DDB2 and Ki67 positive cells in the epidermis and distance to the skin surface (**Figure 5A and 5B**); and (2) distance to nearest blood vessel— showing the distance of immune cells to the nearest endothelial cell in the dermis region (**Figure 5C and 5D**). For illustration purposes, two cases of mild and marked sun exposure are shown in **Fig. 5A-D:** Region 11 (HuBMAP ID: HBM875.KTPB.893) is from the upper arm of a 41-year-old female with mild sun exposure effects (determined from H&E evaluation); https://juyingnan.github.io/vccf_visualization.io/html/region_11.html and Region 7 (HuBMAP ID: HBM384.NNQH.676) is a biopsy from the lower forearm of a 69-year-old male with marked sun exposure effects <u>ttps://juyingnan.github.io/vccf_visualization.io/html/region_7.html.</u> Noteworthy visible differences between the two donors include markedly higher DDB2 positive cells distributed through 0-200 µm of the epidermis and relatively low counts of p53 and Ki67 positive cells in the mild exposure/younger donor. In contrast, the marked exposure/older donor had more p53 and Ki67 positive cells, and markedly lower DDB2 positive cells distributed more within 100-200 µm of the epidermis. There were similar distribution of immune cell counts within 100 µm of the nearest endothelial cell in both donors. Example multiplexed regions of interest and distance overlay for both donors are shown in **Supp. Fig. S11**.

**Figure 5A and 5B:**
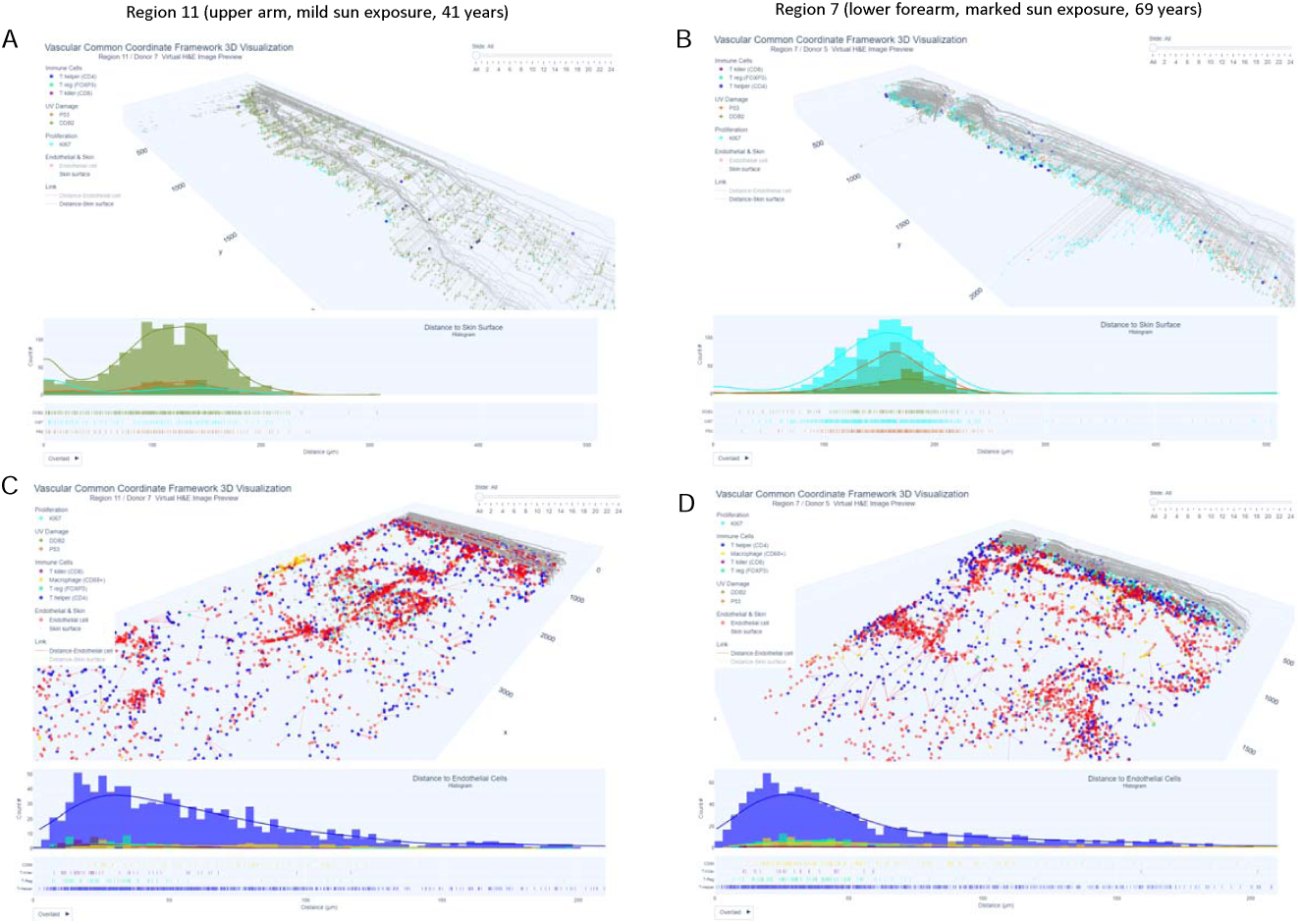
Markers of UV damage, repair and proliferation in epidermis and distance to skin surface represented as 3D distance plots from the skin surface and histogram distribution. **Figure 5C and D** Immune cells and endothelial cells distance to skin surface represented as 3D distance plots from nearest endothelial cells and histogram distribution.

## Discussion

We have presented novel methods for spatially registering data in three dimensions using the HuBMAP registration user interface^13^; selecting antibodies aligned with the anatomical structures, cell types, and biomarker (ASCT+B) tables in support of high quality, ontology aligned data generation; generating 3D volumes of digital skin biopsy data using multiplexed imaging of sequential sections; computing distance distributions of cell damage and proliferation markers to the skin surface; and compiling distance distributions of major immune cells to the nearest blood vessel in support of a vasculature-based human common coordinate framework^36^. Interactive data visualizations allow users to explore spatial patterns of cell type distance distributions in relation to vasculature and position within the epidermis. This 3D workflow is extendable to other cell types (and organs) and will provide a powerful approach for cellular resolution 3D spatial analysis and for constructing a human reference system. All datasets and code are freely available at <u>GitHub - hubmapconsortium/MATRICS-A: Multiplexed Image Three-D Reconstruction and Integrated Cell Spatial -Analysis</u> and via a HuBMAP collection at <u>Samples | HuBMAP</u> (hubmapconsortium.org)

Although 3D reconstruction of multiplexed serial sections is a relatively time-consuming process, it has a significant advantage of generating significantly more cellular data per sample with high resolution. A number of other methods have been used to investigate 3D volumes of organs^12^, however, challenges include antibody penetration (requiring long incubation times), preservation of antigens, tissue architecture distortion, and limitations in the delineation of cell-to-cell relationships on structures spanning several millimeters. Wang et al.^40^ used confocal microscopy to demonstrate lymphatic and blood vessel networks in the human dermis using immunostaining for CD31, Podoplanin, and LYVE-1. Light-sheet microscopy has been used to image skin structure in 3D^41^, but the use of low NA objectives and low magnification to provide a wide field of view results in poor spatial resolution at cellular level. Three-dimensional reconstruction of serially sectioned H&E-stained skin samples has shown the variation in dermis structure and other macro features using the CODA method^42^. Although we did not incorporate reconstruction of H&E histological images, this workflow would be possible using the virtual H&E images generated from the autofluorescence images acquired in this study and would be another informative way to visualize the cellular data. The addition of this dataset is planned for future work.

This study included a small sample of skin specimens sampled across various body locations to account for diversity in anatomical organization and degrees of UV exposure. Acute UV exposure dampens aspects of the immune response, which has positive effects for autoimmune disease, but can impair the response to neoplastic cells^43^. The long-term effects of chronic UV exposure on immune composition have not been well studied and one study has shown restoration of most immune markers to baseline after 14 days^44^. In our study, aging and sun exposure effects were highly correlated, with younger donors more likely to have mild sun exposure effects. While significant differences were not found in immune cell counts by age and sun exposure, more T killer cells were found within 100 μm of endothelial cells in younger patients and this warrants further exploration. An additional consideration when studying skin and interpreting changes in cell counts is anatomical location, which significantly influences the thickness of keratin layer, epidermis and dermis, as well as the distribution and density of adnexal structures such as hair follicles, sebaceous, apocrine and sweat glands etc. ^45,46^. Skin from the face, neck, scalp, and back of hand are significantly exposed to the sun when compared to sites such as trunk, medial aspect of extremities (inner thigh, arm etc.,) and plantar foot (sole), leading to marked differences in the amount of cumulative UV radiation exposure. Partially addressing this, our samples were collected from across the anatomy, including arms, legs, abdomen, and scalp and normalized for total volume, epidermis volume and endothelial cell count to account for sample-to-sample differences. Although we did not find significant differences in p53, our results suggest that DNA repair is more effective in younger/mild sun exposed patients and decreases with age and/or sun exposure. Pilkington et al.^47^ have reported that senescent phenotypes in aged skin can result in reduced skin barrier function and promote a chronic low-level inflammation or “inflammaging” in response to stresses such as repeated UV exposure over the course of chronological aging. Related to this, future areas of research should incorporate more donors and expand racial diversity. Additional insights would be gleaned from incorporation of more aging, senescence, immune (e.g. Langheran cells) and functional immune markers (e.g., exhaustion or activation^48^). It would also be of tremendous value to combine spatial transcriptomic data to further interrogate underlying biology. Importantly, this 3D reconstruction workflow can be applied to any tissue or organ type using any multiplexing technology using sequential section imaging for elucidation of spatial relationships between cell types and vasculature.

## Methods

### Patient samples

Skin biopsies were collected from 12 donors ranging from 32-72 years with a mix of typically UV-exposed and non-exposed anatomical regions (**Supplementary (Supp.) Table S1**). The biopsies were trimmed to size, ranging from 14×12 mm to 47×21 mm in dimension, and embedded in a single block that underwent micro CT imaging. The blocks were then sectioned into 100 5-micron serial sections, numbered in sequence, of which up to 26 of the highest quality in serial succession were selected for further analysis (slide layout shown with virtual H&Es –comprised of pseudo-colored autofluorescence and DAPI^49^ in **Supp. Fig. S2**). All 12 biopsies were spatially registered using the HuBMAP Registration User Interface, and submission of corresponding metadata (donor information, including health status and sample processing) for each sample (https://hubmapconsortium.github.io/ccf-ui/rui/ - **Supp. Fig. S3**). Of the 12 samples, 10 were down selected for further analysis. The two excluded samples included a donor with a benign cyst, but with extensive inflammation and immune cell infiltration compared to other samples. The second sample had a scar which also altered the normal organization of the epidermis and dermis layers. All donors were in good health and cancer free at the time of sample collection, with two donors having chronic diseases (RA and HIV), which is noted in the patient summary table.

### Pathologist review

Virtual H&E images (which are pseudo colored autofluorescence and DAPI images^49^) from each donor were assessed by a pathologist for histopathological changes related to chronic sun exposure such as keratinocytic atypia in the epidermis and solar elastosis changes in the dermis. Accordingly, the specimens were categorized into groups of skin with mild, moderate, and marked sun exposure effects (**Supp. Table S1**). Donors in the mild sun exposure were significantly younger than the moderate-marked exposure donors (42.4 *vs*. 62.2 years, p = 0.008). All virtual H&E images are located at: <u>vccf-visualization-release/vheimages at main · hubmapconsortium/vccf-visualization-release · GitHub</u>

### Micro CT imaging of skin blocks

A Phoenix micro CT system (GE, Wunstorf, Germany) with up to 300 kV/500W was used to generate high resolution CT images of the skin samples. Phoenix micro CT scanners have a high dynamic DXR digital detector array and can produce isotropic images of 1 μm and are frequently used for industrial process control as well as for scientific research applications. Due to its dual tube configuration, detailed 3D information for an extremely wide sample range can be provided. For our purpose a current of 200 kV was found to be optimum in terms of signal-to-noise ratio to generate high quality volumetric isotropic images of 0.016 mm resolution for the embedded skin samples and this also allowed imaging of 12 samples in one block within 30 minutes. Example images for micro CT with corresponding histological section are shown in **Supp. Fig. S3**. The DICOM header with imaging settings is shown in **Supp. Table S2**.

### Antibody validation and multiplexed analysis

All antibodies used in this study were subjected to a standardized characterization process using a tissue microarray (TMA) and appropriate controls to evaluate the specificity and sensitivity of the primary antibody and its dye-conjugated derivative, including the cyclic testing of the dye inactivation treatment compared to single staining^50^. **Supp. Table S3** shows the antibody clones and conjugates used in the study. The 18-marker panel provided coverage for 9 cell types: epithelial, fibroblast, immune cells (macrophage, T helper, T killer, T reg), nerve, myoepithelial, and endothelial cells. These are also highlighted using the HuBMAP <u>A</u>natomical <u>S</u>tructures <u>C</u>ell <u>T</u>ype and <u>B</u>iomarkers (ASCT+B) reporter comparison feature https://hubmapconsortium.github.io/ccf-releases/v1.0/docs/asct-b/skin.html. An example sample region of the skin ASCT+B reporter is shown in **Supp. Fig. S4**. Multiplexed immunofluorescence (MxIF) staining of the skin samples was performed as previously described ^49^ using Cell DIVE™ technology (Leica Microsystems, Issaquah, WA). After de-paraffinization and a two-step antigen retrieval, the FFPE slides were stained with DAPI and imaged in all channels of interest to acquire background autofluorescence (AF) of the tissue. This was followed by primary/secondary and/or direct conjugate antibody staining of up to 3 markers per round plus DAPI, dye deactivation, and repeat staining to collect images of all planned biomarkers. Multiplexed images were automatically registered and processed for illumination correction and autofluorescence subtraction. Each sample was comprised of ∼40 20x fields of view and 20-26 sections were imaged for 3D reconstruction. Example virtual H&E and multiplexed images for two contrasting regions (mild vs. marked sun exposure) are shown in **Supp. Fig. S5**.

### Cell Segmentation and Classification

**Figure 2** and **Supp. Fig. S6** summarize our segmentation model framework. First, an encoder-decoder based deep learning (DL) model^4^ was trained on a small sample (194 DAPI image patches) of manually annotated nuclei from the DAPI images. Multiscale Laplacian of Gaussian (LOG) was introduced along with DAPI images as separate channels to our encoder-decoder based DL model. The LOG feature detects blob like structures in the DAPI images corresponding to the nuclei shape and boundary, thereby providing contextual information to our DL model. Use of multiple channels allows us to train an accurate DL model from a small sample of manually annotated DAPI images. Depth of our encoder-decoder DL model was set to 4 and binary cross entropy was used as a loss function. An unsupervised GMM^5^ was used on cell biomarkers for an automatic probabilistic segmentation of immune cell types (T killer (CD8), T reg (FOXP3), T helper (CD4), macrophages (CD68), endothelial cells (CD31), as well as markers of proliferation (Ki67) and DNA damage (p53) and DNA repair (DDB2) (**Supp. Fig. S7**). Union of probabilities obtained from the GMM model and nuclei segmentation of our DL model was then fused for cell segmentation. Probability values were used to automatically scale (between 0-1) and quantify cells with high expression and determine percentage overlap for a single cell with segmented nuclei. Low percentage overlap was further used to remove imaging artifacts, debris, and cells with low expression. DAPI images were normalized to zero mean unit variance values inside our deep learning framework prior to nuclei segmentation. Secondly, the individual whole slides were normalized between zero mean and unit standard deviation before estimation of biomarker probabilities. Whole slide image specific normalization ensures all images were scaled relative to their intensity distribution and reduces intensity variability often observed between serial whole slide images. The combination of these two normalizations ensured a single cell was segmented based on relative intensity difference between the biomarker and the background and not on absolute intensity distribution that may vary from one slide to another adversely affecting segmentation accuracy. The same GMM was used to automatically segment contiguous structures, independent of nuclei, such as blood vessels (based on CD31 staining) and epithelial masks (based on cytokeratin staining). In such a scenario, we depend on probabilities as obtained from our GMM and automatically threshold our probability using Otsu filters^51^ on the probability map of the biomarkers. **Fig. 2A** shows examples of a multiplexed region of interest from donor 9; epidermis (**Fig. 2B**); cell classification (in this example, p53 positive cells) (**Fig. 2C**); and (**Fig. 2D**) endothelial cells. For validation, a total of 2722 positive and negative cell markers were manually annotated using the annotation function in QuPATH^52^ as follows: CD3: 408; CD4: 281; CD8: 347; FOXP3: 360; CD68: 391, CD31: 352; Ki67: 150; p53: 164; DDB2: 162 and these were used to calculate cell classification sensitivity, specificity and accuracy (**Fig. 2E**). Code can be found at: <u>GitHub - hubmapconsortium/MATRICS-A: Multiplexed Image Three-D Reconstruction and Integrated Cell Spatial - Analysis</u>

### 3D Reconstruction

For 3D volume reconstruction of the autofluorescence (AF) images, a reference AF image was selected for registration of all remaining serial section slides and generation of a 3D tissue block (**Figure 3 A and B**). AF images were automatically segmented using Otsu thresholding, morphological closing, and retaining largest component. AF images were masked prior to affine, and B-spline based deformable registration between the reference image and the serial sections to improve registration accuracy. A block matching strategy was adopted to determine the transformation parameters for the affine registration from masked AF images (**Figure 3C**). The similarity between a block from the reference AF was computed relative to the AF serial sections. The best corresponding block defined the displacement vector for the affine transformation^53^. Normalized cross-correlation similarity was used to determine the block correspondences. Initial affine registration on block correspondences was followed by deformable B-spline based registration^54^ between the reference AF image and the serial section to account for deformation of the serial section during the staining process (**Figure 3C**). Normalized mutual information was chosen as the similarity matrix and maximized to achieve the registration. The transformation map obtained from the registration of the AF images was applied to individual biomarkers for all serial sections to create a 3D volume of endothelial, T killer, T reg, T helper cells and macrophages (**Figure 3D**) and overlaid on 3D AF volume as shown in **Figure 3E**. Slides were randomly picked from each of the skin volumes and segmented biomarkers were overlaid on the biomarker images for a visual validation. Further, density of cell types was verified from one slide to another in the 3D volume using the visualization generated from the 3D VCCF model. To resolve any issues dealing with higher or lower signal intensity for a slide, any discrepancies in cell density of a biomarker were identified and a higher or lower probability threshold was used on the probabilistic segmentation to achieve a better segmentation. Such manual biomarker probability adjustments were necessary in less than 2% of the whole slide images. Code can be found at: <u>GitHub - hubmapconsortium/MATRICS-A: Multiplexed Image Three-D Reconstruction and Integrated Cell Spatial -Analysis</u>

### Cell data aggregation and analytics

For patient level comparison, cell counts within the regions of interest (either entire sample or the epidermis region in 3D reconstructed data) were aggregated at patient level by each cell type. Then the quantities were normalized by the volume of the region of interest. To create the epidermis volume, epidermis segmented from each whole slide image was given the same transformation as the AF volume to create a 3D reconstructed volume of the epidermis. Number of voxels of the 3D volume were automatically determined using the Insight Toolkit (ITK) software^22^. Based on the quantities aggregated at donor level, statistical hypothesis tests were performed to understand the correlation between the normalized cell count vs. age/UV-exposure. To measure the correlation with age, Spearman’s correlation was quantified and tested. For UV exposure (mild vs. moderate-marked), Wilcoxon-test was performed. For both statistical tests, Benjamini & Hochberg’s multiple testing correction was applied^55^.

### Distance metrics

Two types of distances were computed based on the segmentation result: (1) the distance between the centroid of the nucleus for immune cells (macrophage, T helper, T killer, T reg) and the edge of the nearest blood vessel (endothelial cells) and (2) the distance between the centroid of cells with UV damage markers (DDB2, p53) and proliferation markers (KI67) to the edge of the skin surface. To speed up distance calculations for the 13,489 immune cells and 12,407 damage/proliferation markers across the ten 3D regions, a filter was applied to calculate the square root of the distance for only those cells/markers that fell within the range of the current minimum value of the distance (i.e., cells/markers whose distance on either the x, y, or z axis is greater than the current minimum distance are not included in the distance computation). This approach considerably reduced run time and memory load. Code can be found at:

### Violin plots of cell counts and distance from endothelial cells or from cell surface

Violin plots in **Fig. 4** combine a box plot and a density plot to display the probability density of the data at different values. The interquartile range is represented by the box in the center, and the extended line above/below the box shows the upper (max) and lower (min) adjacent values. In the online interactive version, a kernel density estimation is shown on each side of the box to show the distribution shape of the distance from the cell to the nearest blood vessel. The median value of the distance is represented by the line dot in the middle of the box, and the mean value is represented by the dash line. This was used to depict the distribution of immune cells (macrophages (CD68+), T helper (CD3+, CD4+), and regulatory T reg cells (CD3+, CD4+, FOXP3+) and T killer cells (CD3+, CD8+), as well as the sum of all four) and distances from endothelial cells in regions with marked sun exposure and mild sun exposure for all donors https://juyingnan.github.io/vccf_visualization.io/html/violin_cell_all_region.html and spanning the age range of donors https://juyingnan.github.io/vccf_visualization.io/html/violin_cell.html. Similar plots were generated for distance of epithelial cells positive for Ki67, p53 and DDB2 across the entire sample area and within the epidermis, for all donors and separated by age https://juyingnan.github.io/vccf_visualization.io/html/epidermis/violin_damage_all_region_epidermis.html; https://juyingnan.github.io/vccf_visualization.io/html/epidermis/violin_damage_epidermis.html

### Interactive visualization of cell and marker distances

Interactive 3D visualizations of cell and biomarker distances were implemented using the Plotly 3D visualization package. For Ki67, p53 and DDB2 markers within the epidermis and distance to skin surface: https://juyingnan.github.io/vccf_visualization.io/html/epidermis; for immune cell distances to blood vessels in the dermis: https://juyingnan.github.io/vccf_visualization.io. The distance calculation result was visualized in two ways: (1) a 3D view projecting all immune cell nuclei, damage/proliferation markers, blood vessels, and their shortest distances (lines) in tissue space and (2) 2D histograms showing the distribution of distances between nuclei and blood vessels, and between damage/proliferation markers and the skin surface. The visualization of distance links has been optimized by adding invisible links that unite all of the existing links into a single polyline which significantly reduces the size of the vector data and memory usage, allowing for responsive online interaction with about 20,000 nodes in a web browser. In the 3D view, each cell nucleus is rendered as a small circle in the 3D visualization, whereas each UV damage/proliferation marker is rendered as a cross (see legend for cell and biomarker type colors). Blood vessels and epithelial masks (based on positive cytokeratin staining) are rendered as a collection of red circles. The slider in the top-right corner of each 3D view allows the user to view each tissue layer separately. The histogram views provide information about the distribution of distances for further analysis. The short lines beneath the histograms indicate the relationship between all the samples and the histogram bars. The histogram can be displayed in three different layouts: Overlaid (by default), Stacked, or Grouped, see selection button on lower left.

The visualizations can be exported as an HTML file for online presentation and exploration, or as a vector image for static viewing.

