## Supplementary Figures for "Human Digital Twin: Automated Cell Type Distance Computation and 3D Atlas Construction in Multiplexed Skin Biopsies"

### Slide 1
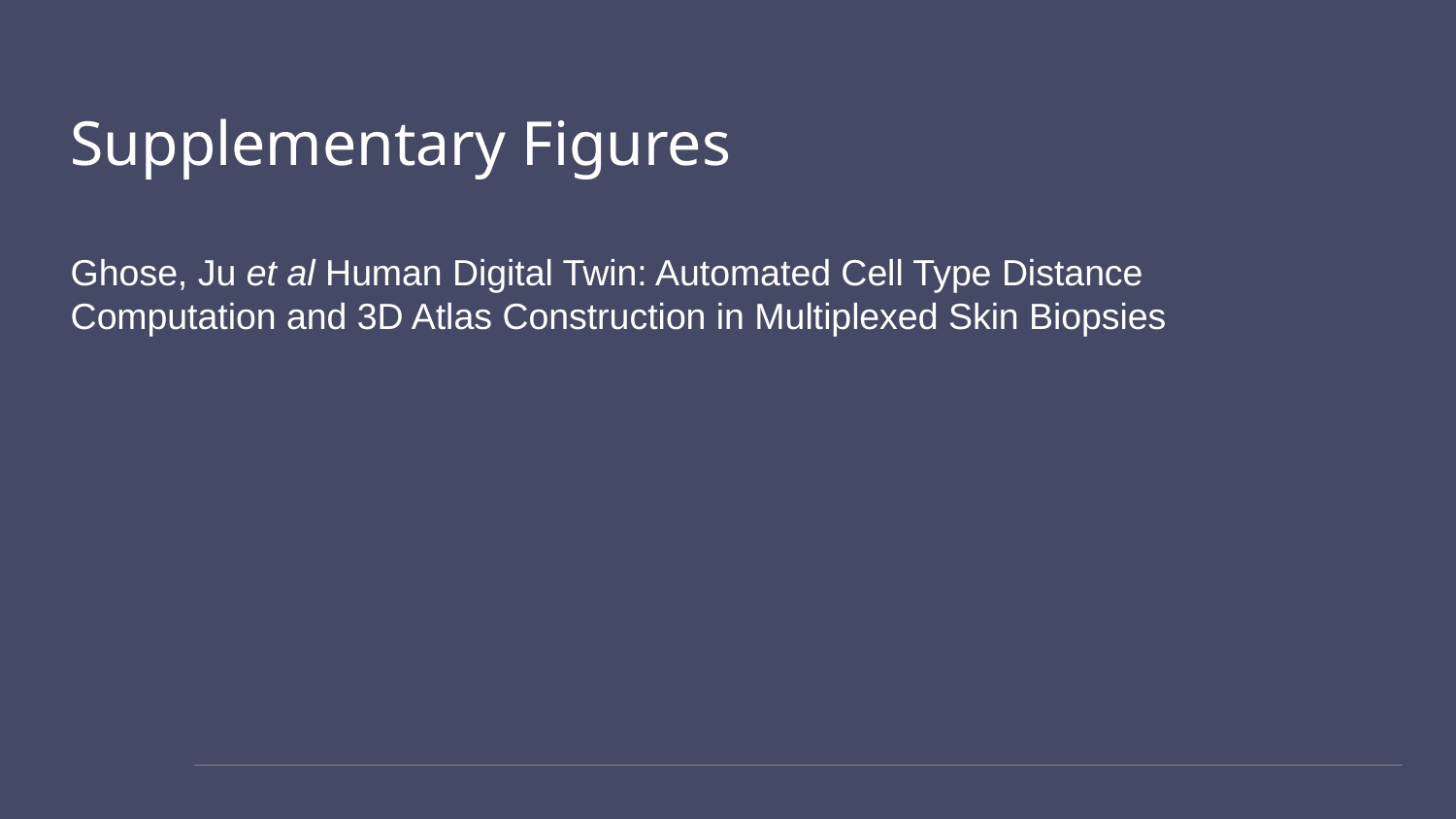

#### Supplementary Figures Ghose, Ju et al Human Digital Twin: Automated Cell Type Distance Computation and 3D Atlas Construction in Multiplexed Skin Biopsies

### Slide 2
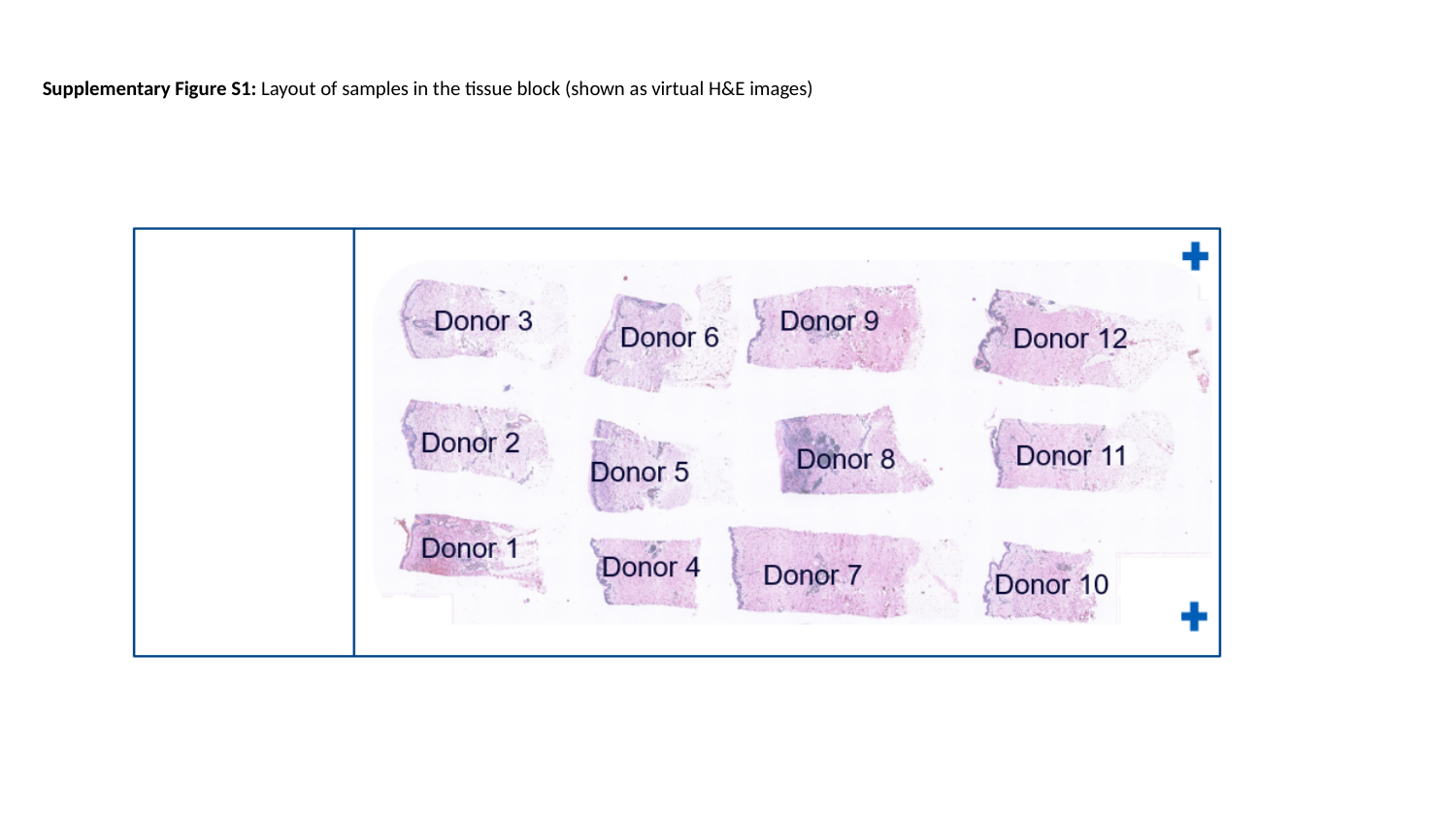

Supplementary Figure S1: Layout of samples in the tissue block (shown as virtual H&E images)

### Slide 3
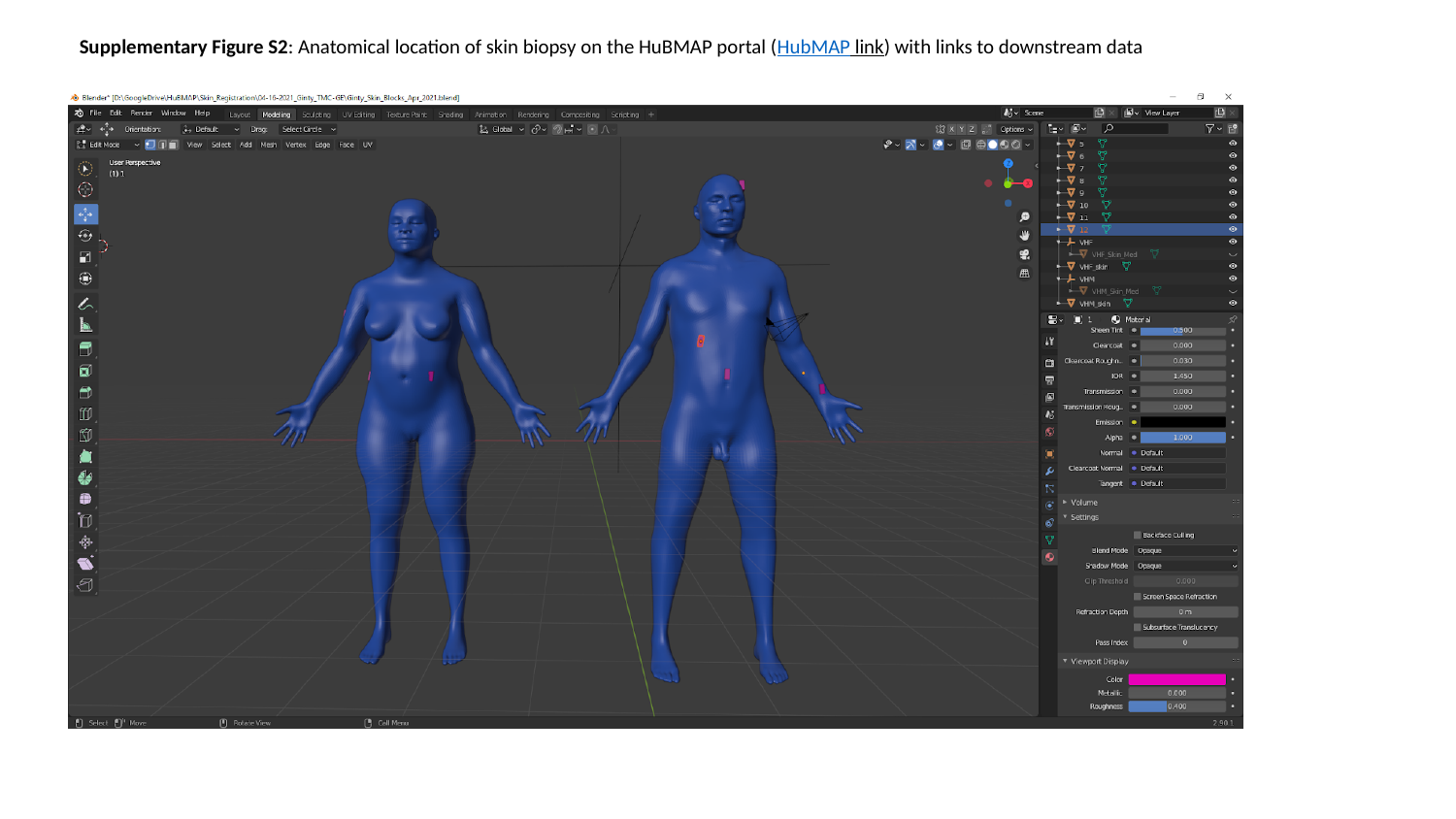

Supplementary Figure S2: Anatomical location of skin biopsy on the HuBMAP portal (HubMAP link) with links to downstream data

### Slide 4
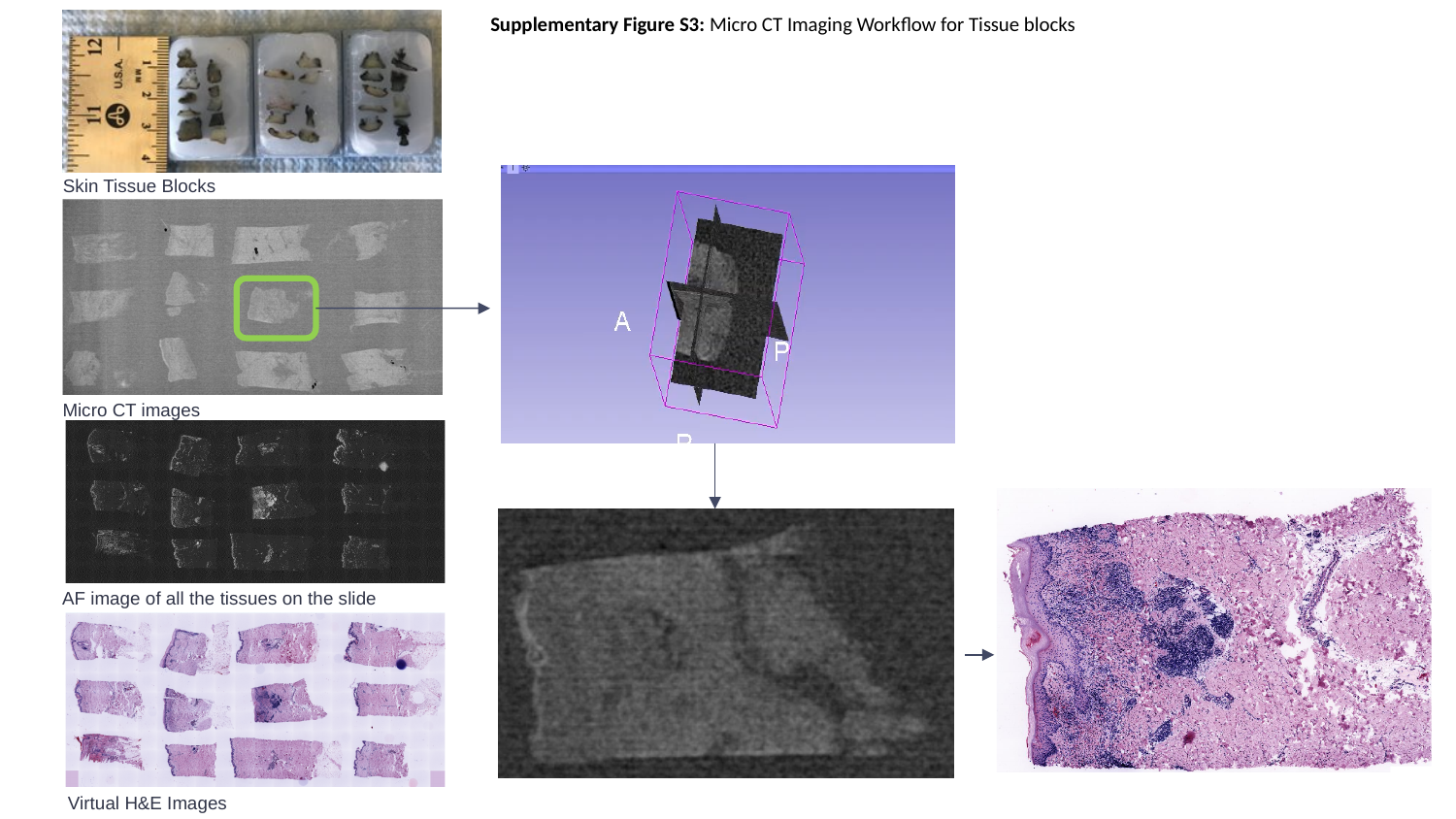

Supplementary Figure S3: Micro CT Imaging Workflow for Tissue blocks
Skin Tissue Blocks
Micro CT images
AF image of all the tissues on the slide
4
Virtual H&E Images

### Slide 5
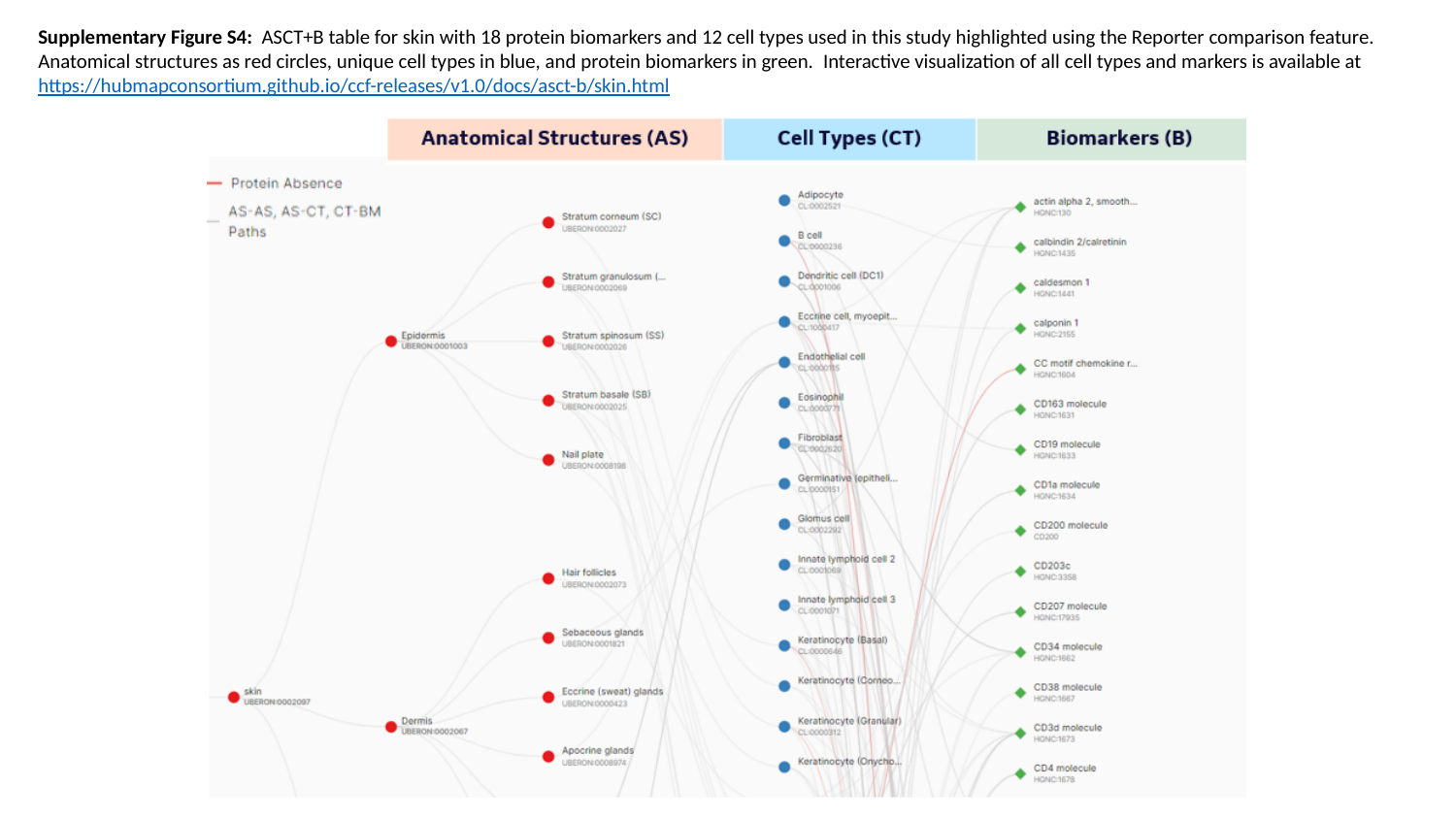

Supplementary Figure S4:  ASCT+B table for skin with 18 protein biomarkers and 12 cell types used in this study highlighted using the Reporter comparison feature. Anatomical structures as red circles, unique cell types in blue, and protein biomarkers in green.  Interactive visualization of all cell types and markers is available at https://hubmapconsortium.github.io/ccf-releases/v1.0/docs/asct-b/skin.html

### Slide 6
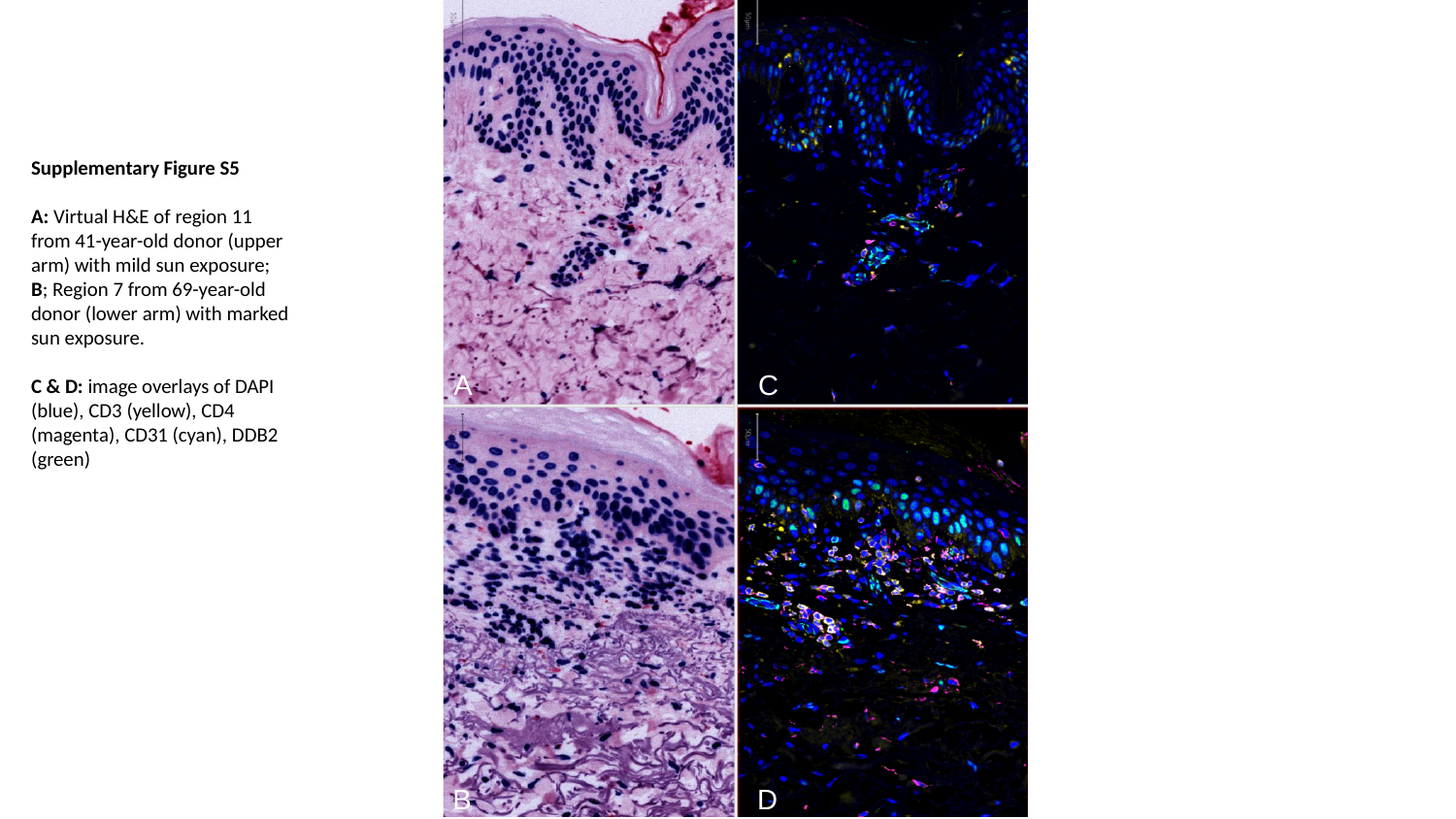

Supplementary Figure S5
A: Virtual H&E of region 11 from 41-year-old donor (upper arm) with mild sun exposure; B; Region 7 from 69-year-old donor (lower arm) with marked sun exposure.
C & D: image overlays of DAPI (blue), CD3 (yellow), CD4 (magenta), CD31 (cyan), DDB2 (green)
A
C
D
B

### Slide 7
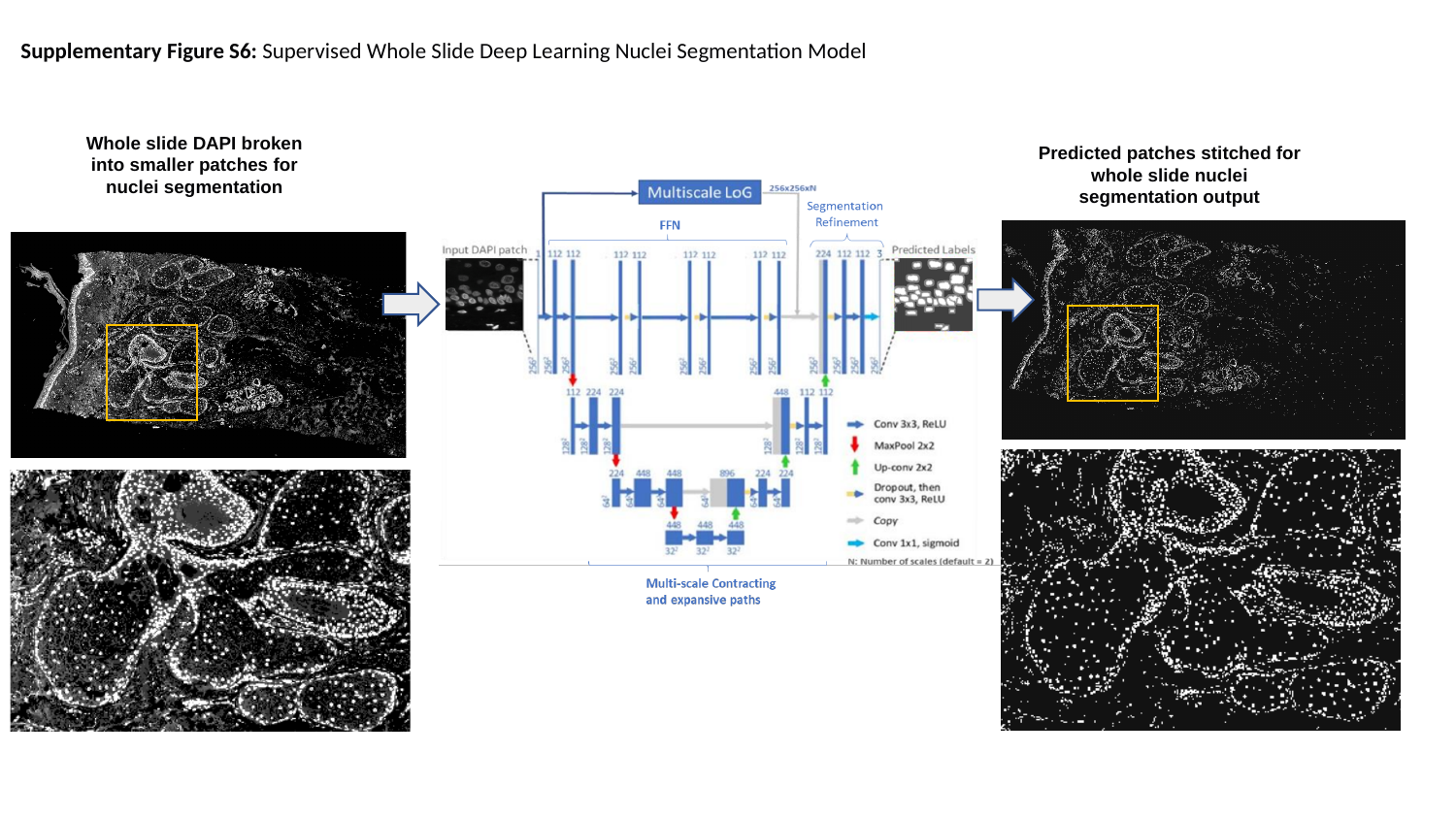

Supplementary Figure S6: Supervised Whole Slide Deep Learning Nuclei Segmentation Model
Whole slide DAPI broken into smaller patches for nuclei segmentation
Predicted patches stitched for whole slide nuclei segmentation output

### Slide 8
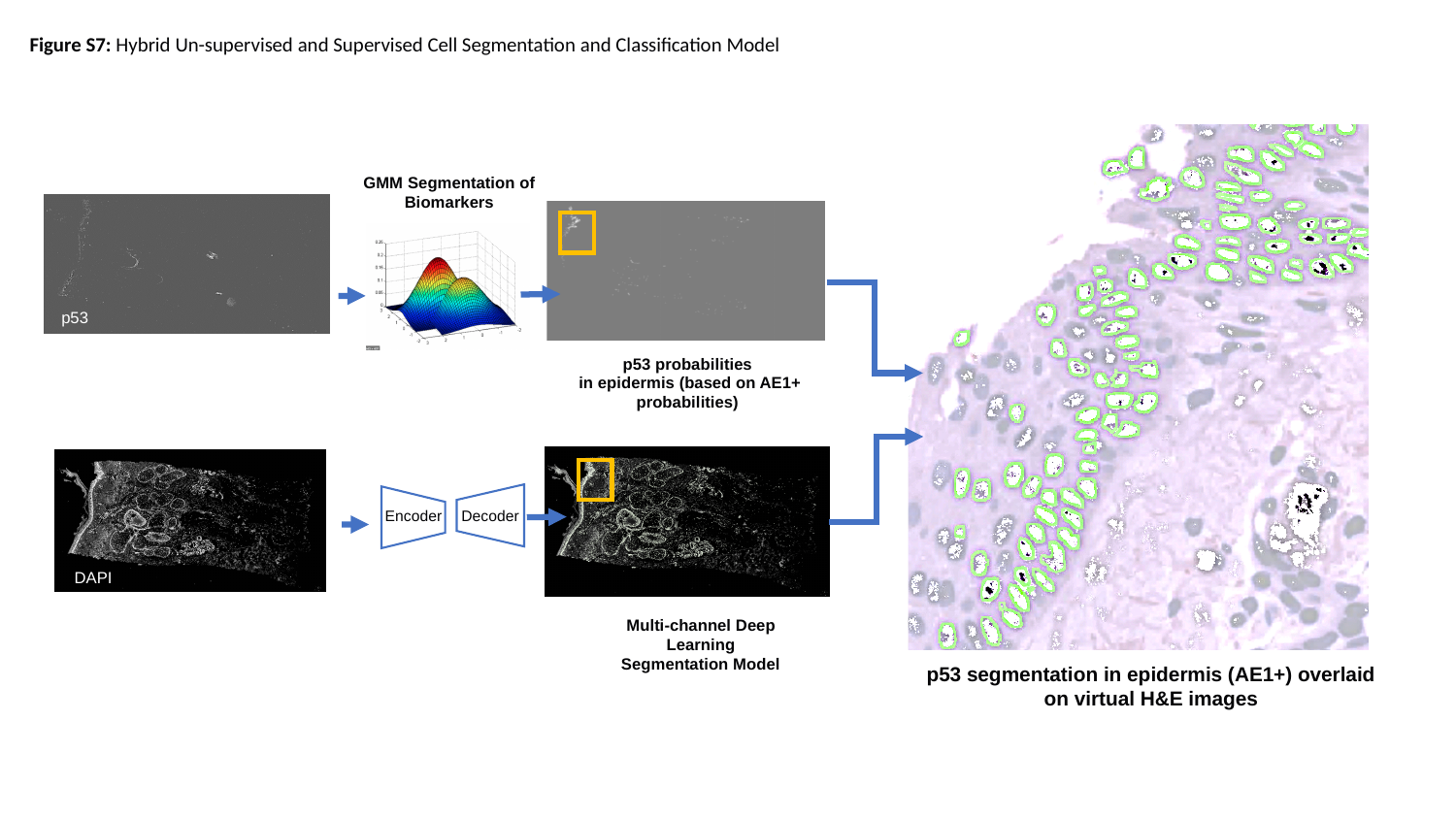

Figure S7: Hybrid Un-supervised and Supervised Cell Segmentation and Classification Model
Decoder
p53 probabilities
in epidermis (based on AE1+ probabilities)
Encoder
Multi-channel Deep Learning Segmentation Model
p53 segmentation in epidermis (AE1+) overlaid on virtual H&E images
GMM Segmentation of Biomarkers
p53
p53
DAPI

### Slide 9
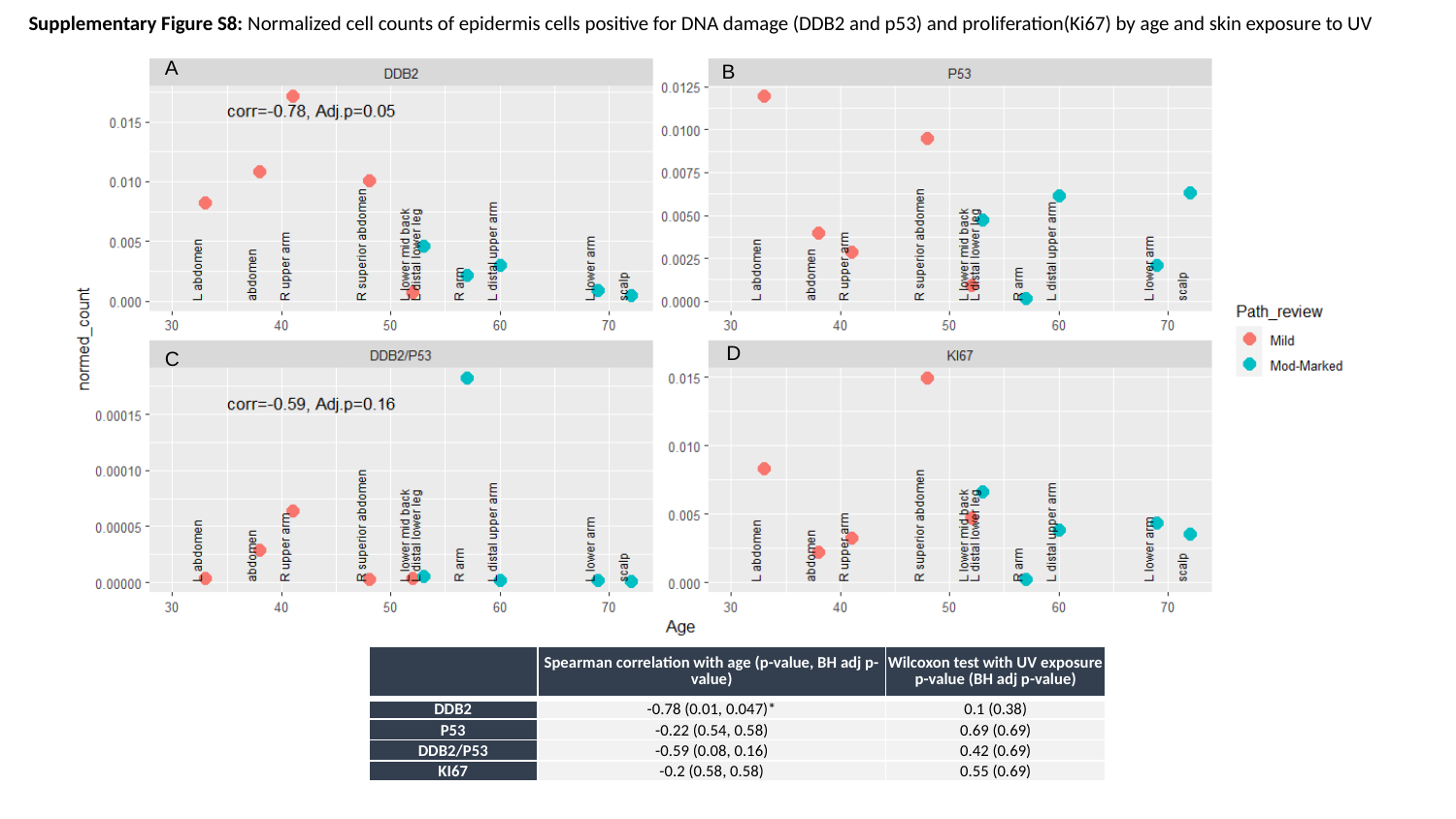

Supplementary Figure S8: Normalized cell counts of epidermis cells positive for DNA damage (DDB2 and p53) and proliferation(Ki67) by age and skin exposure to UV
A
B
D
C
| | Spearman correlation with age (p-value, BH adj p-value) | Wilcoxon test with UV exposure p-value (BH adj p-value) |
| --- | --- | --- |
| DDB2 | -0.78 (0.01, 0.047)\* | 0.1 (0.38) |
| P53 | -0.22 (0.54, 0.58) | 0.69 (0.69) |
| DDB2/P53 | -0.59 (0.08, 0.16) | 0.42 (0.69) |
| KI67 | -0.2 (0.58, 0.58) | 0.55 (0.69) |

### Slide 10
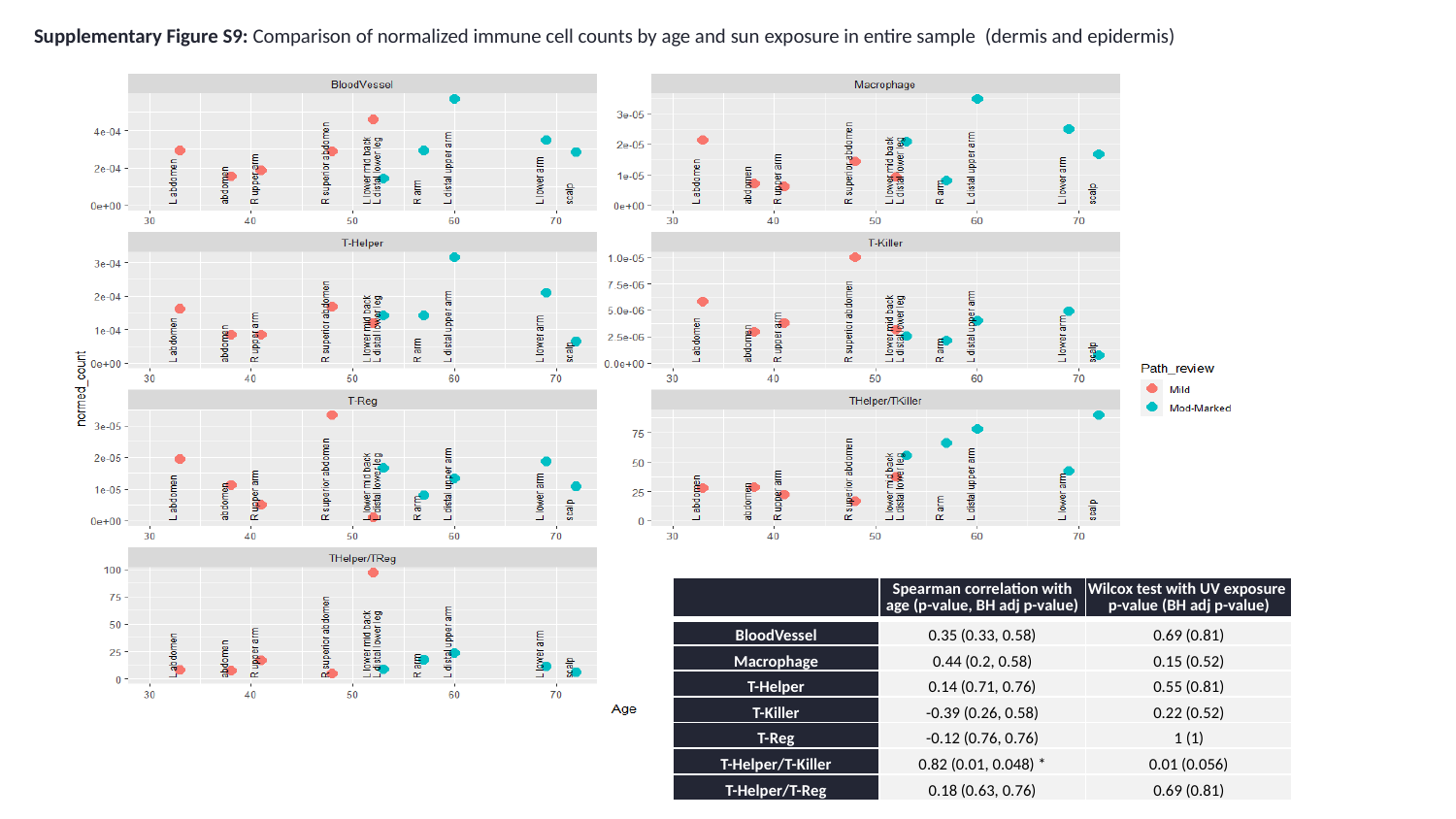

Supplementary Figure S9: Comparison of normalized immune cell counts by age and sun exposure in entire sample  (dermis and epidermis)
| | Spearman correlation with age (p-value, BH adj p-value) | Wilcox test with UV exposure p-value (BH adj p-value) |
| --- | --- | --- |
| BloodVessel | 0.35 (0.33, 0.58) | 0.69 (0.81) |
| Macrophage | 0.44 (0.2, 0.58) | 0.15 (0.52) |
| T-Helper | 0.14 (0.71, 0.76) | 0.55 (0.81) |
| T-Killer | -0.39 (0.26, 0.58) | 0.22 (0.52) |
| T-Reg | -0.12 (0.76, 0.76) | 1 (1) |
| T-Helper/T-Killer | 0.82 (0.01, 0.048) \* | 0.01 (0.056) |
| T-Helper/T-Reg | 0.18 (0.63, 0.76) | 0.69 (0.81) |

### Slide 11
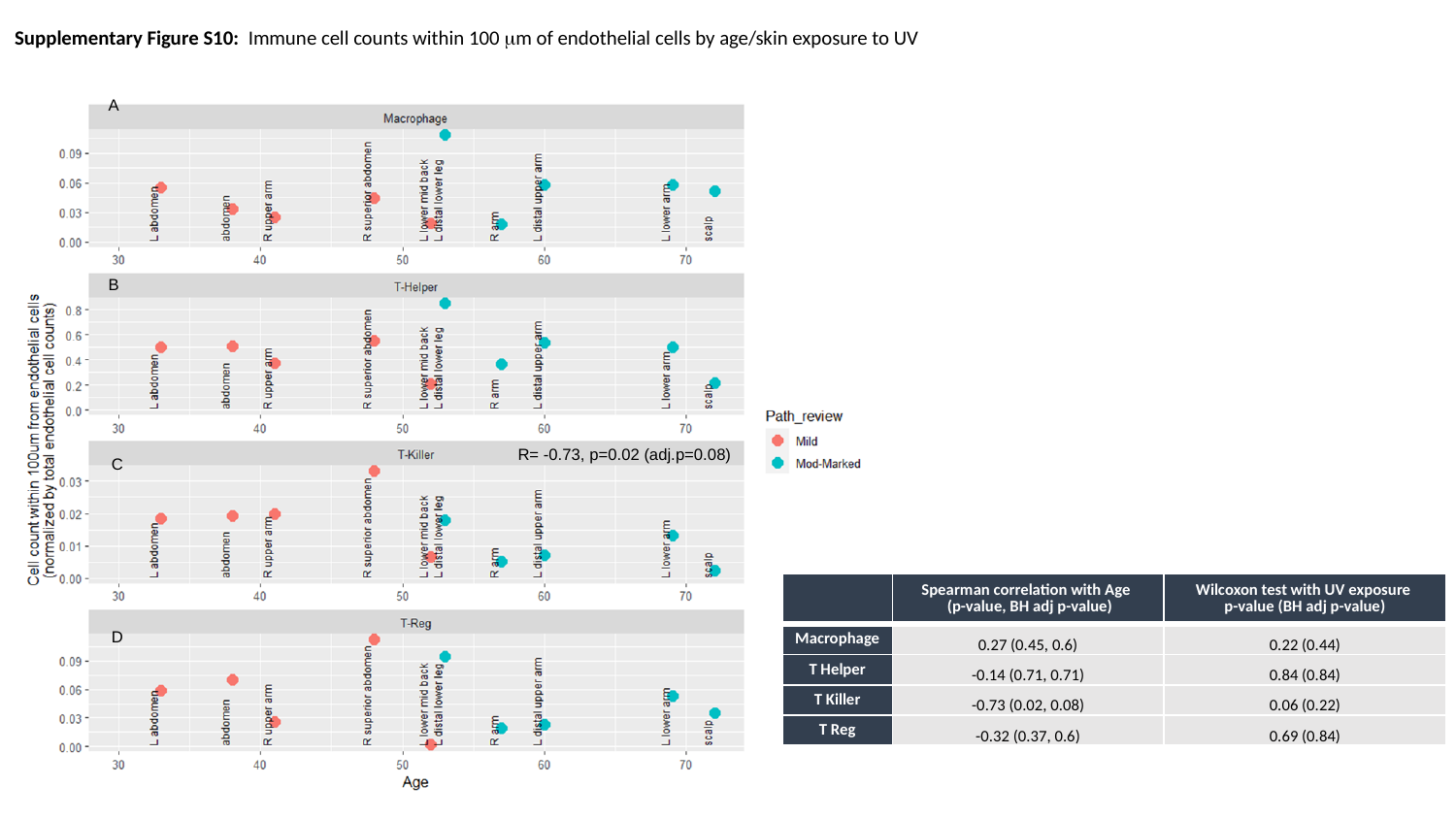

Supplementary Figure S10: Immune cell counts within 100 μm of endothelial cells by age/skin exposure to UV
A
B
R= -0.73, p=0.02 (adj.p=0.08)
C
| | Spearman correlation with Age   (p-value, BH adj p-value) | Wilcoxon test with UV exposure  p-value (BH adj p-value) |
| --- | --- | --- |
| Macrophage | 0.27 (0.45, 0.6) | 0.22 (0.44) |
| T Helper | -0.14 (0.71, 0.71) | 0.84 (0.84) |
| T Killer | -0.73 (0.02, 0.08) | 0.06 (0.22) |
| T Reg | -0.32 (0.37, 0.6) | 0.69 (0.84) |
D

### Slide 12
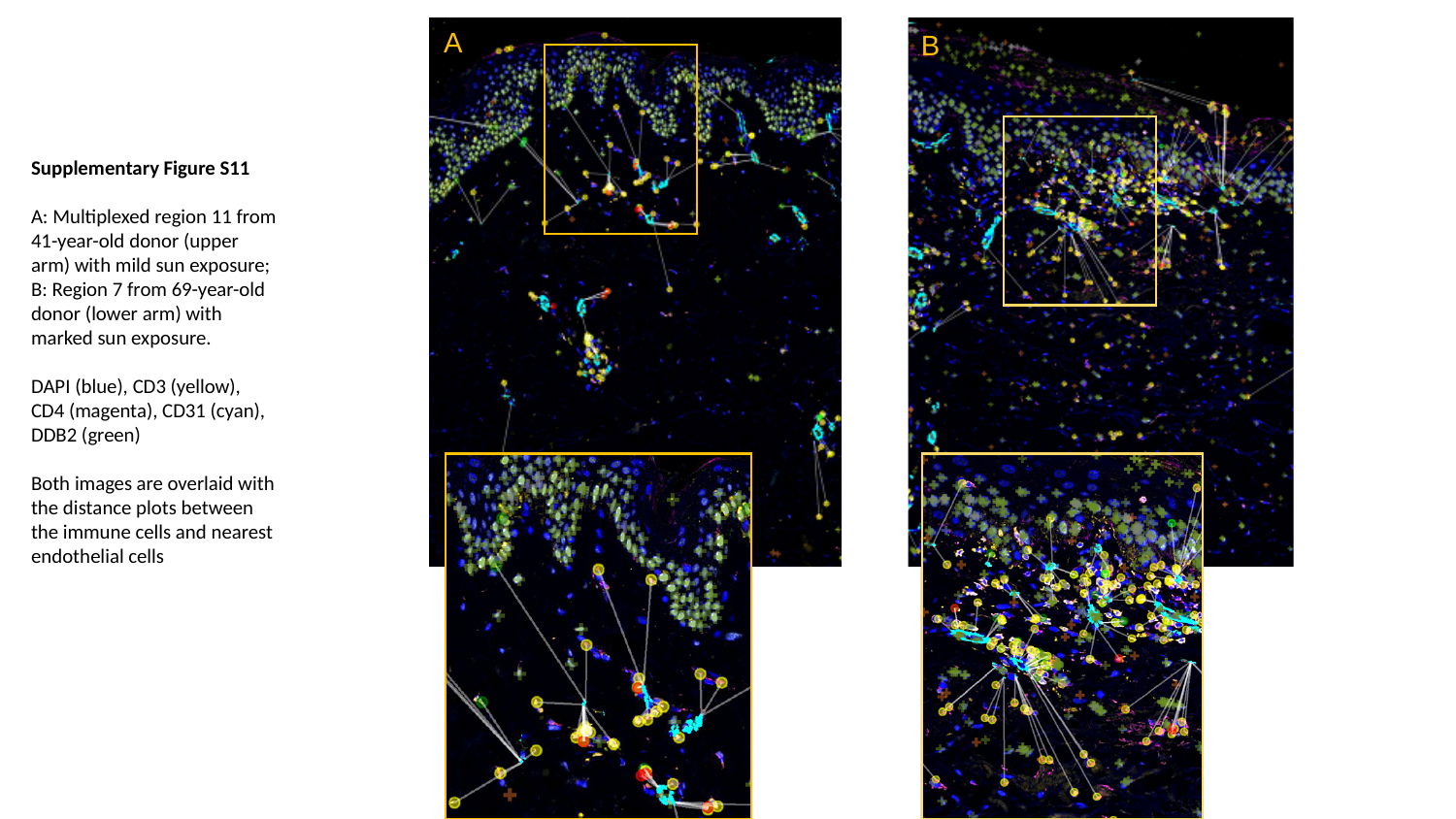

A
B
Supplementary Figure S11
A: Multiplexed region 11 from 41-year-old donor (upper arm) with mild sun exposure; B: Region 7 from 69-year-old donor (lower arm) with marked sun exposure.
DAPI (blue), CD3 (yellow), CD4 (magenta), CD31 (cyan), DDB2 (green)
Both images are overlaid with the distance plots between the immune cells and nearest endothelial cells
