## Supplementary Tables for "Human Digital Twin: Automated Cell Type Distance Computation and 3D Atlas Construction in Multiplexed Skin Biopsies"

### Slide 1
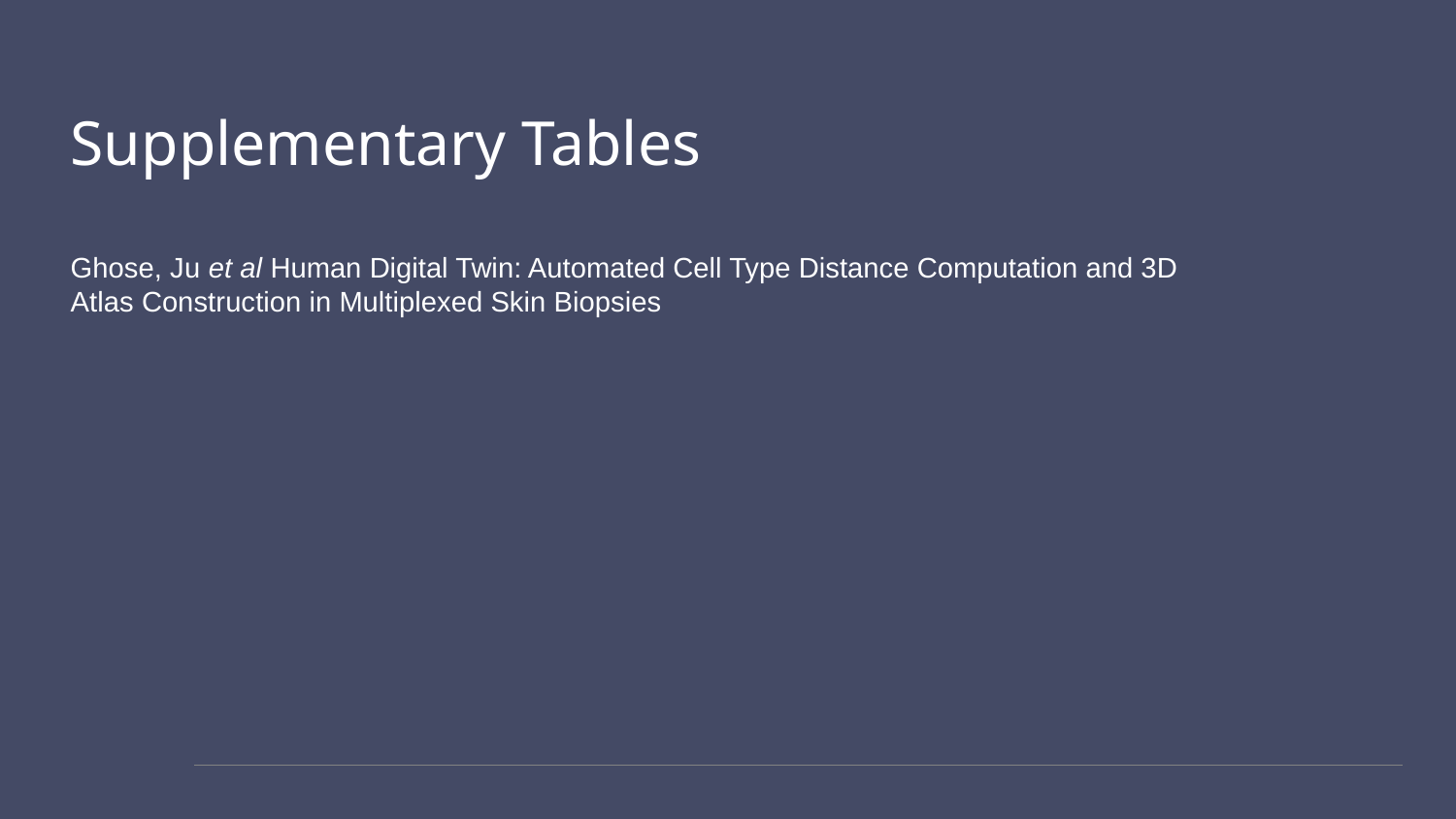

#### Supplementary Tables Ghose, Ju et al Human Digital Twin: Automated Cell Type Distance Computation and 3D Atlas Construction in Multiplexed Skin Biopsies

### Slide 2
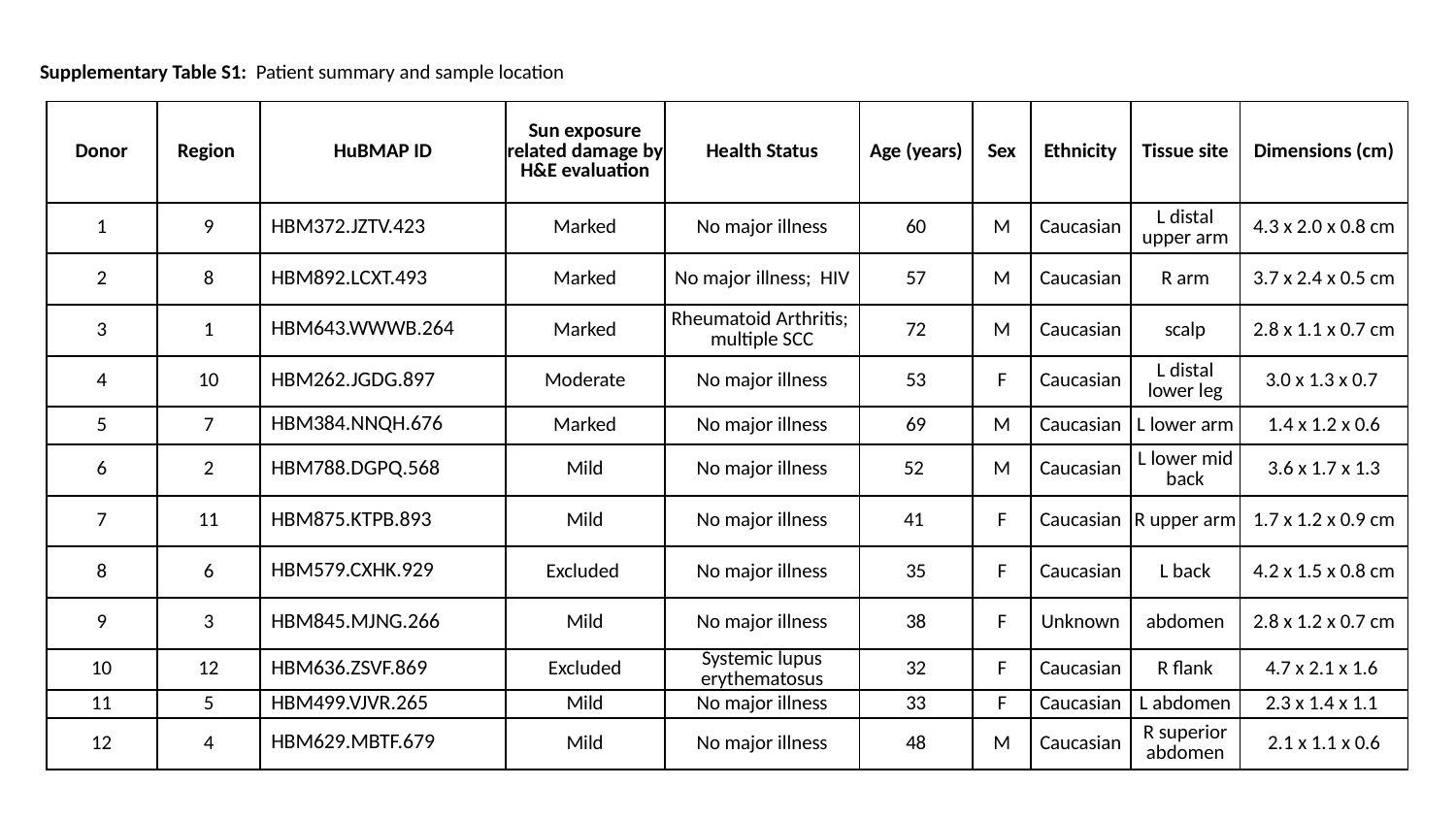

Supplementary Table S1: Patient summary and sample location
| Donor | Region | HuBMAP ID | Sun exposure related damage by H&E evaluation | Health Status | Age (years) | Sex | Ethnicity | Tissue site | Dimensions (cm) |
| --- | --- | --- | --- | --- | --- | --- | --- | --- | --- |
| 1 | 9 | HBM372.JZTV.423 | Marked | No major illness | 60 | M | Caucasian | L distal upper arm | 4.3 x 2.0 x 0.8 cm |
| 2 | 8 | HBM892.LCXT.493 | Marked | No major illness;  HIV | 57 | M | Caucasian | R arm | 3.7 x 2.4 x 0.5 cm |
| 3 | 1 | HBM643.WWWB.264 | Marked | Rheumatoid Arthritis; multiple SCC | 72 | M | Caucasian | scalp | 2.8 x 1.1 x 0.7 cm |
| 4 | 10 | HBM262.JGDG.897 | Moderate | No major illness | 53 | F | Caucasian | L distal lower leg | 3.0 x 1.3 x 0.7 |
| 5 | 7 | HBM384.NNQH.676 | Marked | No major illness | 69 | M | Caucasian | L lower arm | 1.4 x 1.2 x 0.6 |
| 6 | 2 | HBM788.DGPQ.568 | Mild | No major illness | 52 | M | Caucasian | L lower mid back | 3.6 x 1.7 x 1.3 |
| 7 | 11 | HBM875.KTPB.893 | Mild | No major illness | 41 | F | Caucasian | R upper arm | 1.7 x 1.2 x 0.9 cm |
| 8 | 6 | HBM579.CXHK.929 | Excluded | No major illness | 35 | F | Caucasian | L back | 4.2 x 1.5 x 0.8 cm |
| 9 | 3 | HBM845.MJNG.266 | Mild | No major illness | 38 | F | Unknown | abdomen | 2.8 x 1.2 x 0.7 cm |
| 10 | 12 | HBM636.ZSVF.869 | Excluded | Systemic lupus erythematosus | 32 | F | Caucasian | R flank | 4.7 x 2.1 x 1.6 |
| 11 | 5 | HBM499.VJVR.265 | Mild | No major illness | 33 | F | Caucasian | L abdomen | 2.3 x 1.4 x 1.1 |
| 12 | 4 | HBM629.MBTF.679 | Mild | No major illness | 48 | M | Caucasian | R superior abdomen | 2.1 x 1.1 x 0.6 |

### Slide 3
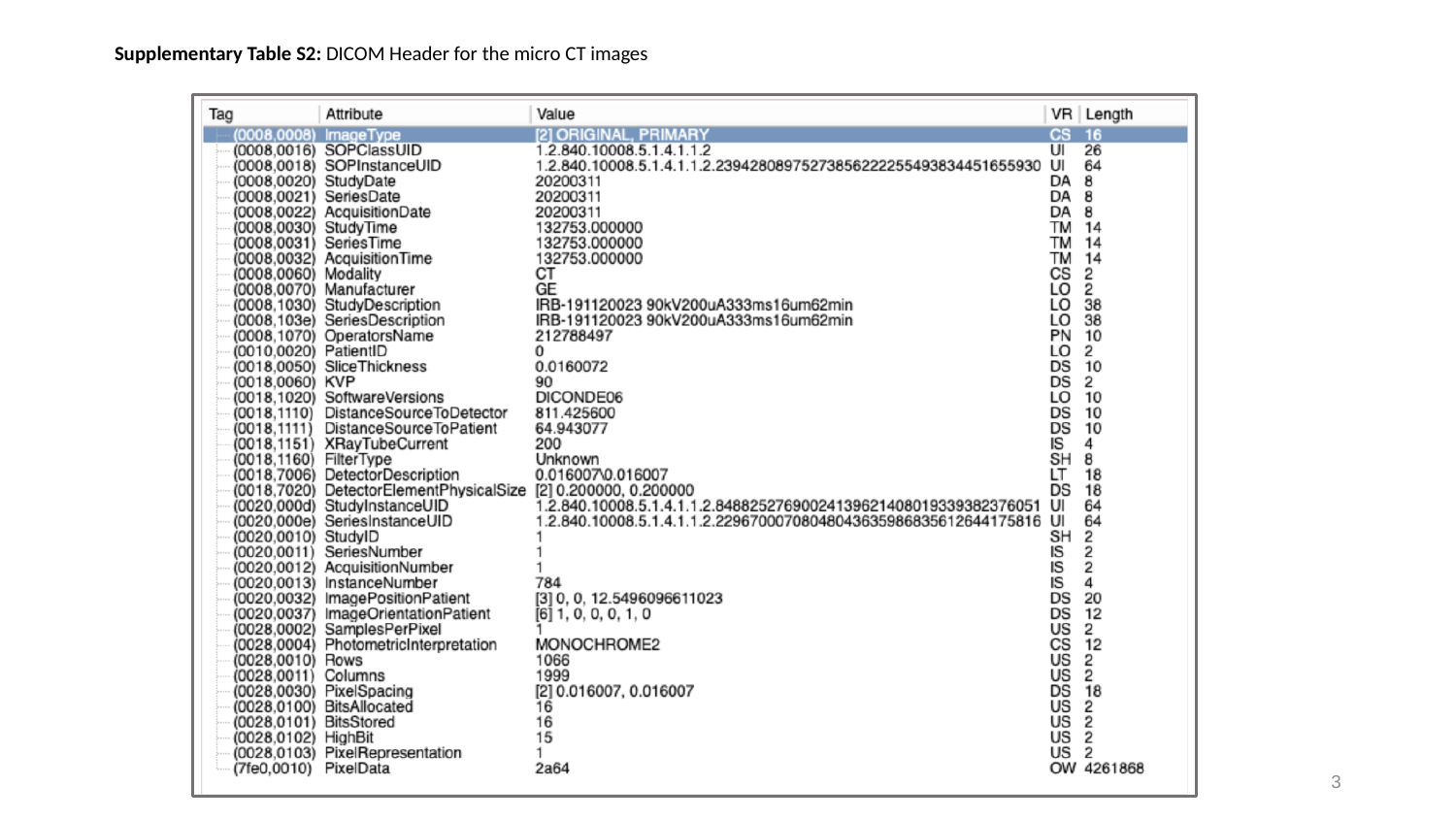

Supplementary Table S2: DICOM Header for the micro CT images
3

### Slide 4
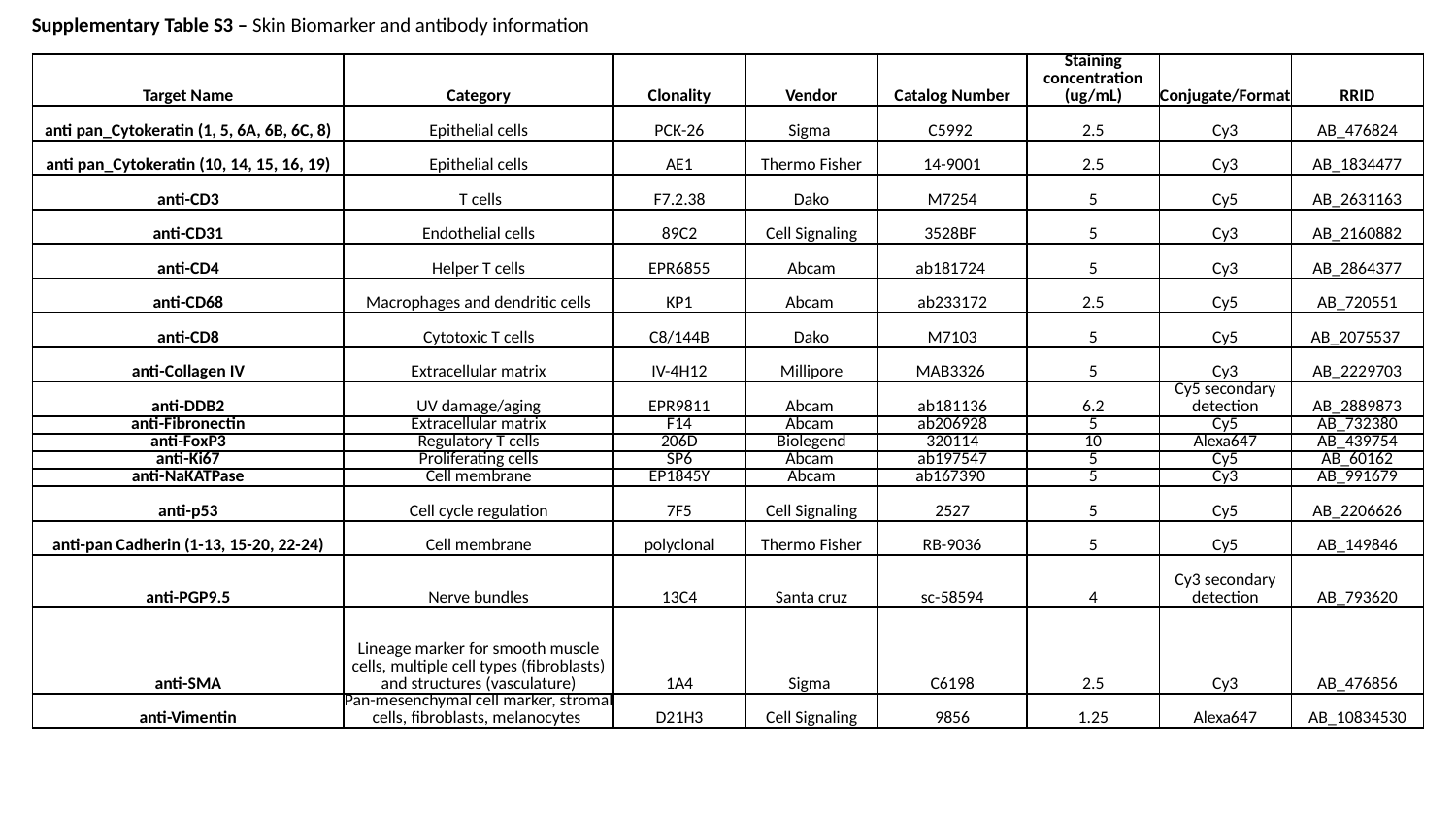

Supplementary Table S3 – Skin Biomarker and antibody information
| Target Name | Category | Clonality | Vendor | Catalog Number | Staining concentration (ug/mL) | Conjugate/Format | RRID |
| --- | --- | --- | --- | --- | --- | --- | --- |
| anti pan\_Cytokeratin (1, 5, 6A, 6B, 6C, 8) | Epithelial cells | PCK-26 | Sigma | C5992 | 2.5 | Cy3 | AB\_476824 |
| anti pan\_Cytokeratin (10, 14, 15, 16, 19) | Epithelial cells | AE1 | Thermo Fisher | 14-9001 | 2.5 | Cy3 | AB\_1834477 |
| anti-CD3 | T cells | F7.2.38 | Dako | M7254 | 5 | Cy5 | AB\_2631163 |
| anti-CD31 | Endothelial cells | 89C2 | Cell Signaling | 3528BF | 5 | Cy3 | AB\_2160882 |
| anti-CD4 | Helper T cells | EPR6855 | Abcam | ab181724 | 5 | Cy3 | AB\_2864377 |
| anti-CD68 | Macrophages and dendritic cells | KP1 | Abcam | ab233172 | 2.5 | Cy5 | AB\_720551 |
| anti-CD8 | Cytotoxic T cells | C8/144B | Dako | M7103 | 5 | Cy5 | AB\_2075537 |
| anti-Collagen IV | Extracellular matrix | IV-4H12 | Millipore | MAB3326 | 5 | Cy3 | AB\_2229703 |
| anti-DDB2 | UV damage/aging | EPR9811 | Abcam | ab181136 | 6.2 | Cy5 secondary detection | AB\_2889873 |
| anti-Fibronectin | Extracellular matrix | F14 | Abcam | ab206928 | 5 | Cy5 | AB\_732380 |
| anti-FoxP3 | Regulatory T cells | 206D | Biolegend | 320114 | 10 | Alexa647 | AB\_439754 |
| anti-Ki67 | Proliferating cells | SP6 | Abcam | ab197547 | 5 | Cy5 | AB\_60162 |
| anti-NaKATPase | Cell membrane | EP1845Y | Abcam | ab167390 | 5 | Cy3 | AB\_991679 |
| anti-p53 | Cell cycle regulation | 7F5 | Cell Signaling | 2527 | 5 | Cy5 | AB\_2206626 |
| anti-pan Cadherin (1-13, 15-20, 22-24) | Cell membrane | polyclonal | Thermo Fisher | RB-9036 | 5 | Cy5 | AB\_149846 |
| anti-PGP9.5 | Nerve bundles | 13C4 | Santa cruz | sc-58594 | 4 | Cy3 secondary detection | AB\_793620 |
| anti-SMA | Lineage marker for smooth muscle cells, multiple cell types (fibroblasts) and structures (vasculature) | 1A4 | Sigma | C6198 | 2.5 | Cy3 | AB\_476856 |
| anti-Vimentin | Pan-mesenchymal cell marker, stromal cells, fibroblasts, melanocytes | D21H3 | Cell Signaling | 9856 | 1.25 | Alexa647 | AB\_10834530 |
